# Long-read transcriptome assembly reveals vast transcriptional complexity in the placenta associated with metabolic and endocrine function

**DOI:** 10.1101/2025.06.26.661362

**Authors:** Sean T. Bresnahan, Hannah Yong, William H. Wu, Sierra Lopez, Jerry Kok Yen Chan, Frédérique White, Pierre-Étienne Jacques, Marie-France Hivert, Shiao-Yng Chan, Michael I. Love, Jonathan Y. Huang, Arjun Bhattacharya

## Abstract

The placenta is critical for fetal development and mediates the effects of pregnancy complications on offspring metabolic health, yet it is often poorly characterized in genomic studies. Existing transcriptomic analyses rely on adult tissue-based references, which overlook developmentally important isoform diversity. We used largest-in-class long-read RNA-seq (N=72) to create a comprehensive placental transcriptome reference, identifying 37,661 high-confidence isoforms (14,985 novel) across 12,302 genes (2,759 novel). Contrary to characterizations of the placenta as a “transcriptomic void,” we found transcriptional breadth and complexity comparable to adult tissues, with extraordinary splicing diversity in genes controlling obesity, lactogen production and growth, including 108 distinct *CSH1* (placental lactogen) isoforms. This improved reference offers two advantages: First, it reduced inferential uncertainty in isoform quantification by 30% and increased the yield of high-confidence transcripts. Applying this reference to short-read RNA-seq datasets (N=344) of gestational diabetes mellitus (GDM), we found that placental transcription mediated 36% of GDM effects on birth weight, with novel *CSH1* isoforms identified as key mediators. We further uncovered ancestry-specific effects, with distinct *CSH1* isoforms mediating larger effects in European (24.4%) than Asian (13.4%) populations. Our results establish that utilizing long-read-based, tissue-specific transcriptomic annotations is critical, enabling isoform-resolved analyses that provide greater sensitivity than conventional gene-level approaches for understanding placental function and context-specific variation across diverse biobanks.

## Introduction

The placenta, a transient organ unique to pregnancy, serves as the critical interface between maternal and fetal environments. Beyond its essential roles in nutrient transfer, gas exchange, and waste elimination, the placenta functions as a master regulator of the intrauterine environment through complex neuroendocrine and immunological signaling pathways that influence fetal development and metabolic programming of later-in-life health outcomes.^1–9^

Gestational diabetes mellitus (GDM), characterized by glucose intolerance with onset during pregnancy, affects approximately 16% of pregnancies worldwide and significantly impacts both maternal and fetal health.^9^ While decreasing insulin sensitivity characterizes physiological pregnancy, GDM involves insulin insufficiency relative to the exaggerated decline in maternal insulin sensitivity and glucose metabolism, potentially through altered placenta-mediated endocrine signaling such as the pro-diabetogenic human placental lactogen (hPL) encoded by *CSH1.*^10^ GDM-exposed fetuses experience placenta-mediated alterations in intrauterine signals promoting macrosomia, altered body composition with increased adiposity, and heightened risk for metabolic disorders later in life.^11,12^ These associations are in line with the Developmental Origins of Health and Disease (DOHaD) hypothesis, which posits that the intrauterine environment shapes developmental trajectories with lasting implications for offspring health.^13–17^

### Limitations of short-read sequencing and current annotations restrict understanding of placental biology

Despite the placenta’s central role in prenatal development, it has been notably underrepresented in large-scale tissue-specific genomic and transcriptomic initiatives such as the Genotype-Tissue Expression Project (GTEx), the NIH Roadmap Epigenomics Consortium, and the Encyclopedia of DNA Elements Consortium (ENCODE).^18–20^ These limited resources have restricted our understanding of placenta-specific gene regulation and transcriptional breadth and complexity relevant to fetal growth and metabolic programming.

Developing tissues, including the placenta, exhibit distinct patterns of alternative splicing and isoform usage compared to adult tissues.^20–23^ Using adult tissue or species-level aggregate transcriptome references like GENCODE^24^ for placental transcriptomics obscures tissue-specific regulatory mechanisms critical to understanding placental function. This limitation is compounded by the inherent constraints of short-read RNA sequencing approaches, which face significant challenges in accurately resolving transcript isoforms due to read-to-transcript mapping ambiguity.^25^ When sequenced fragments are compatible with the exons of multiple isoforms, common for genes with complex splicing patterns, probabilistic read assignment introduces significant uncertainty in estimating isoform-level expression.^26^ This inferential uncertainty is particularly problematic for developing tissues exhibiting extensive alternative splicing in functionally important genes.^22^

Analyses of existing placental RNA-sequencing datasets have focused overwhelmingly on aggregated gene-level measurements,^5,14,21,23,27–34^ missing critical biological information by obscuring isoforms that may be particularly susceptible to genetic or environmental effects relevant to complex traits.^35–41^ This gene-centric view fails to capture scenarios where pathological exposures are mediated through changes in specific isoforms that could be masked or compensated by expression of other, potentially non-functional isoforms, or have opposing functions on the same biological process.^35^ For example, previous studies that challenged the role of classical reproductive hormones in GDM, based on findings that circulating hPL levels do not correlate with insulin sensitivity changes during pregnancy,^42^ measured aggregate hormone levels without distinguishing between functionally distinct isoforms that may have differential effects on maternal metabolism.

### Placental long-read sequencing improves functionally relevant expression annotations

Long-read RNA sequencing technologies generate individual reads of up to 15-20 kilobases, sufficient to capture entire transcripts from end to end, providing direct evidence of exon combinations and revealing comprehensive transcriptional breadth and complexity.^43–47^ Recent studies demonstrate that over half of the isoforms detected using long-read RNA sequencing were missed by traditional short-read approaches in the same tissue samples,^48–50^ highlighting a significant gap in current understanding of transcriptome regulation. Previous work, including from our group, demonstrated that alternative splicing patterns and isoform usage explain substantially more variance in complex phenotypes than gene expression, with modeling of isoforms rather than genes increasing discovery of transcriptome-mediated genetic associations with complex traits by approximately 60%.^35,37,51^

Here, we develop the first comprehensive transcriptome reference for placenta derived from N = 72 long-read RNA-sequencing libraries and orthogonal sequencing data. Our analysis identified 37,661 high-confidence transcripts representing 12,302 genes, including isoforms of both known and potentially novel genes, with pregnancy-related genes exhibiting extraordinary isoform diversity compared to other human tissues. We show that this assembly offers two complementary advantages: (1) improved specificity and accuracy of transcript quantification in short-read datasets by reducing mapping ambiguity and (2) the ability to identify novel isoform-level regulatory mechanisms that may be critical for understanding GDM-associated modulation of fetal development. We leveraged this annotation to study GDM and fetal growth, demonstrating how improved isoform resolution reveals novel molecular mechanisms underlying complex pregnancy outcomes. Our approach uncovered previously uncharacterized placental isoforms of genes involved in glucose metabolism, placental lactogen production and fetal growth regulation that mediate the effects of maternal hyperglycemia on birth weight. These findings demonstrate that long-read sequencing-derived placental transcript annotations are essential for understanding pregnancy pathophysiology and detecting mechanistic signals in observational studies of environmental and intrauterine exposures. Given the improved quantification precision and novel mechanistic insights demonstrated here, our placenta reference transcriptome provides a valuable resource for reanalyzing existing short-read datasets to uncover previously missed isoform-level regulatory mechanisms.

## Results

### Long-read sequencing reveals extensive transcript diversity in human placenta

In two pregnancy follow-up studies,^28,52^ we generated long-read transcriptome data from villous placental tissue of N = 72 term, live births without known placentation pathologies, such as intrauterine growth restriction and pre-eclampsia, including 50% from pregnancies affected by GDM **(Figure 1A)**. Using the Oxford Nanopore platform, we sequenced cDNA libraries obtaining approximately 310.8 million reads. After stringent quality control and alignment to hg38 (Methods), we retained approximately 72 million high-quality reads with an average read length of 1.07 kilobases (kb) (SD = 1.25 kb, range = 0.25–22.2 kb) **(Table S1)**. These reads mapped to 68,002 distinct transcripts with a mean length of 1.36 kb (SD = 1.67 kb, range = 0.06–205 kb) **(Figure S1A)** and an average of 4.5 exons per transcript **(Figure S1B)**. These transcripts represented 23,069 genes, with an average of 2.95 isoforms per gene (SD = 3.84) **(Figure S1C)**. Our analysis captured 20,803 of 63,187 genes annotated in GENCODE v45 and identified 2,266 novel genes including 102 fusion products of multiple genes **(Table S2)**.

**Figure 1.**
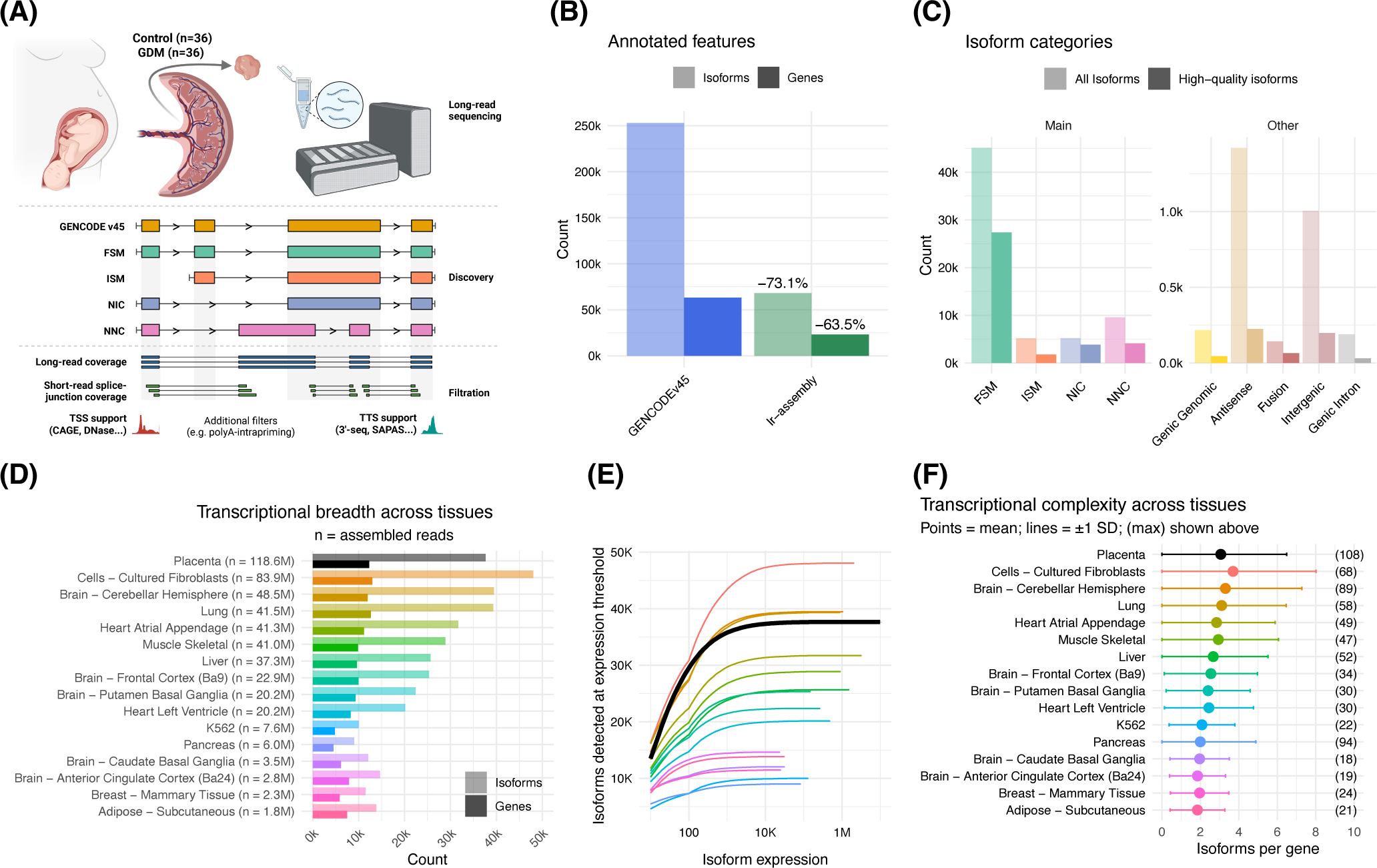
The placenta exhibits transcriptional breadth and complexity comparable to other adult tissues, challenging its characterization as a transcriptomic void. **(A)** Nanopore cDNA libraries were prepared from villous placenta tissue of term, live births from mothers with and without gestational diabetes mellitus (GDM). In the discovery phase, transcripts were assembled and compared to the GENCODE v45 annotations to classify isoforms into structural categories: full splice matches (FSM), incomplete splice matches (ISM), novel in catalog (NIC), novel not in catalog (NNC), and other classes (fusion products and intergenic, genic, intronic and antisense transcripts). In the filtration phase, transcripts were retained if they had strong orthogonal support for long-read coverage, short-read coverage of splice junctions, support for transcription start sites (TSS) and termination sites (TTS) from relevant sequencing modalities (e.g. CAGE for TSS, 3’-seq for TTS), and passed other quality control measures (e.g. no evidence of poly(A)-intrapriming). (B) Genes and transcripts in the GENCODE v45 reference compared to the unfiltered placenta reference transcriptome. (C) Transcripts in the placenta reference transcriptome stratified by structural category. Transparent bars represent the full set of transcripts, while opaque bars indicate the high-confidence transcripts. (D) Transcriptional breadth across tissues, visualized by the count of long-read defined genes and isoforms in placenta compared to 15 tissues or cell lines in GTEx^47^ v9. (E) The number of isoforms detected at increasing read coverage thresholds across tissues (data for placenta shown by thick black line). (F) Transcriptional complexity across tissues, visualized by the mean and standard deviation of isoforms per gene. Maximum values are indicated in brackets.

### Nearly one-quarter of placental transcripts represent novel isoforms

Among transcripts annotated to known genes (n = 65,386 [96.15%]) we identified three major categories by comparison to GENCODE v45: transcripts matching known annotations, novel isoforms of known genes, and transcripts of novel genes **(Figure 1B-C)**. Most transcripts of known genes matched existing annotations, classified as full splice match (FSM: n = 45,082 [68.95%]), where all splice junctions exactly matched a reference transcript, or incomplete splice match (ISM: n = 5,176 [7.91%]), where a subset of splice junctions matched a reference.

We identified 14,957 novel transcripts (22.87% of transcripts expressed in placenta) associated with 5,630 known genes (26.93% of genes expressed in placenta). These novel isoforms averaged 0.79 kb in length (SD = 0.47 kb, range = 0.08–4.4 kb) with mean of 4.5 exons per transcript. The novel transcripts are categorized as: “novel not in catalog” (NNC, n = 9,578 [63.9%]), with at least one previously unannotated splice donor or acceptor site; “novel in catalog” (NIC, n = 5,190, [34.6%]), combining known splice sites in previously unobserved patterns; or “genic” transcripts (n = 217 [1.45%]), which partially mapped to 152 known genes but with insufficient splice junction matches to be classified as NIC or NNC. Additionally, we identified 2,759 transcripts representing novel genes, categorized as: antisense (n = 1,421, [51.5%]), intergenic (n = 1,006 [36.5%]), originating within introns of known genes (n = 189 [6.8%]), or fusion products spanning multiple genes (n = 143 [5.2%]).

### Orthogonal data sources validate high-confidence placental transcript models

For downstream analyses, we filtered out low-confidence isoforms using criteria from previous long-read transcriptome assembly studies (Methods).^36,45–49^ We used orthogonal short-read RNA-seq, DNase I hypersensitivity site sequencing (DNase-seq), Cap Analysis of Gene Expression sequencing (CAGE-seq), and sequencing of alternative polyadenylation sites (SAPAS) data to validate splice junction coverage and transcription start and termination sites (TSS and TTS, respectively) **(Table S16)**. Transcript models showed robust support **(Table S2)** with an average of 554.23 long-reads (Nanopore; SD = 3.89 × 10^4^) **(Figure S2A)**, and an average minimum splice junction support of 2.33 × 10^4^ short-reads (Illumina; SD = 2.47 × 10^5^) **(Figure S2B)**. We observed strong enrichment of TSS near CAGE-seq or DNase-seq peaks (median distance = 90 bp with 23,740 [34.9%] of transcripts located within 50 bp) **(Figure S2C)**. Similarly, TTS showed enrichment near alternative polyadenylation sites **(Figure S2D)** or PolyA motifs **(Figure S2E)** (median distance = 15 bp with 26,549 [40.5%] of transcripts located within 5 bp).

Our quality filtering identified few problematic transcripts: n = 2,243 isoforms (3.3%) showed evidence of nonsense mediated decay (NMD), n = 483 (0.7%) exhibited RT-switching artifacts, and n = 4 (0.005%) showed evidence of PolyA intrapriming **(Figure S2F)**. While NMD transcripts can be biologically meaningful, incomplete assemblies or artifacts can also produce premature termination codons. Therefore, for NMD-predicted transcripts, we required additional evidence of substantial 5’ expression to distinguish genuine NMD substrates from assembly artifacts. Of 2,243 NMD-predicted transcripts in novel categories, only 892 were filtered exclusively due to insufficient 5’ expression evidence; the remaining 1,351 were ultimately filtered for other quality control reasons. After removing 30,341 low-confidence isoforms (44.62%; **Table S3**), our placenta transcript assembly contained 37,661 high-confidence isoforms **(Figure 1C)** with mean length of 1.60 kb (SD = 1.52 kb, range = 0.08–34.63 kb; **Figure S3A**) and an average of 5.62 exons per transcript **(Figure S3B)**. Unless explicitly stated, all subsequent analyses reported here refer to this filtered high confidence set, which represents 12,302 genes (mean = 3.06, SD = 3.43 isoforms per gene), including 11,854 genes in the GENCODE v45 and 448 novel genes **(Figure S3C)**. Leveraging tissue-specific expression data from the Human Protein Atlas,^53^ genes corresponding to the most abundant transcripts in our long-read data were significantly enriched for genes expressed in the placenta and not in other tissues across multiple abundance thresholds **(Table S4; Figure S3D)**. Among the 500 most abundant placental long-read transcripts, we observed a 4.3-fold enrichment for placenta-specific genes (adjusted p = 7.85 × 10^-7^), suggesting our sequencing successfully captured genes with established placenta-restricted expression patterns.

### Novel transcripts show distinct structural characteristics

Compared to transcripts of known genes, transcripts of novel genes were significantly less abundant in the long-reads (Mann-Whitney-Wilcoxon test: W = 1.07 × 10^7^, p < 2.2 × 10^-16^) **(Figure S4A)**, shorter in length (W = 1.31 × 10^7^, p < 2.2 × 10^-16^) **(Figure S4B)**, and contained fewer exons (W = 1.43 × 10^7^, p < 2.2 × 10^-16^) **(Figure S4C)**. Similarly, novel transcripts of annotated genes were significantly less abundant than FSM transcripts (W = 1.25 × 10^8^, p < 2.2 × 10^-16^) and shorter in length (W = 1.58 × 10^8^, p < 2.2 × 10^-16^), although they contained comparable numbers of exons (W = 1.15 × 10^8^, p = 0.99). We identified substantial diversity in the number of multi-exonic RNA isoforms per gene (range: 1–108) with over half of all detected genes (n = 7,343 [59.69%]) expressing more than one isoform **(Figure 2A)**. The number of detected isoforms was correlated with the number of detected exons per isoform (r = 0.87, p < 2.2 × 10^-16^) and the strength of this correlation was stronger among highly expressed genes (r = 0.93, p < 2.2 × 10^-16^), likely reflecting greater sensitivity for detecting splicing events from abundantly expressed genes **(Figure S4D)**.

**Figure 2.**
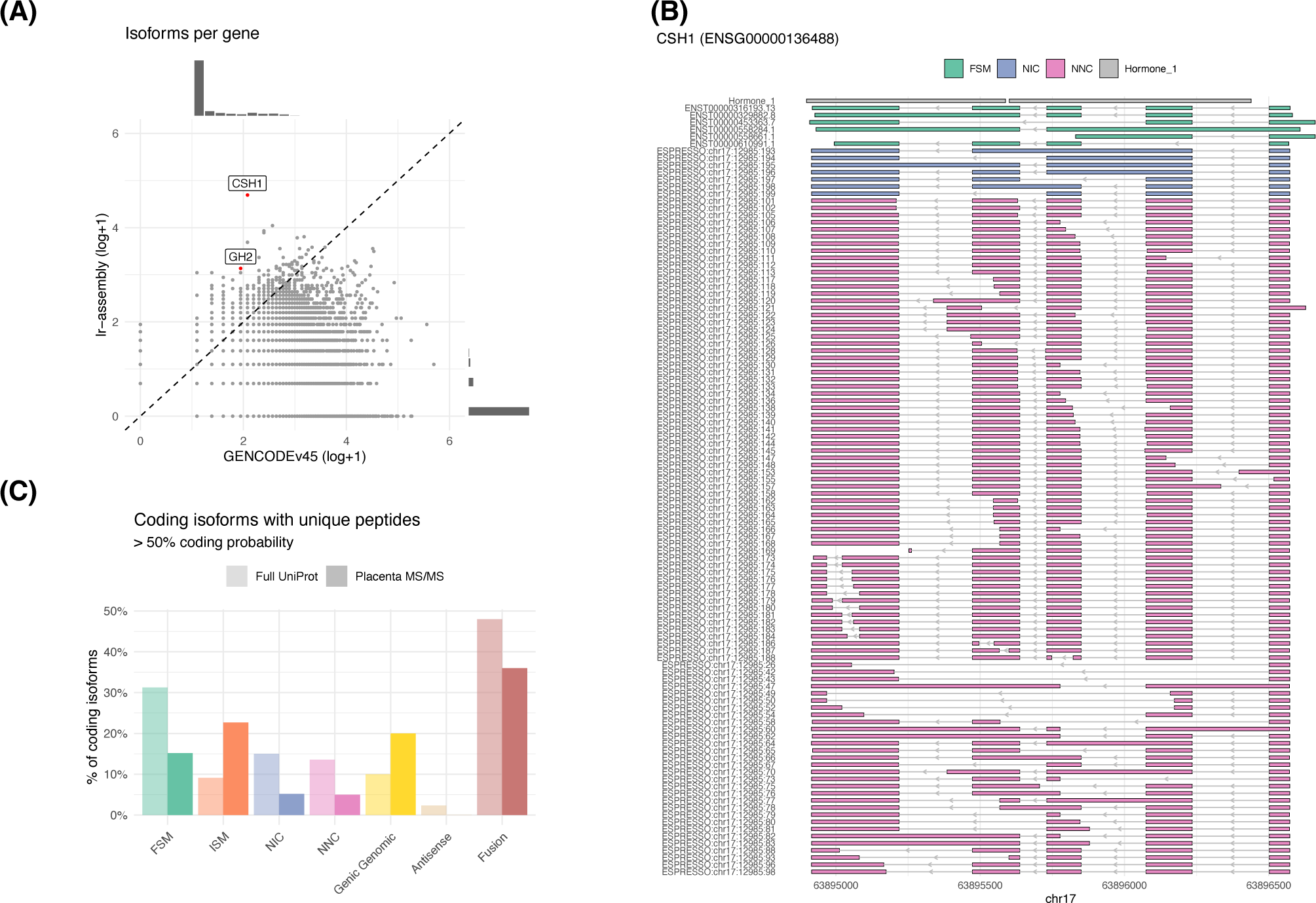
The placenta exhibits extraordinary splicing diversity of pregnancy-related genes. **(A)** Isoforms per gene in the GENCODE v45 reference compared to the high-confidence placenta reference transcriptome (lr-assembly). **(B)** High-confidence transcripts of *CSH1* expressed in the placenta, including a protein domain track for Hormone 1 indicated in grey. **(C)** Transcripts with predicted open reading frames (ORFs) and support for protein coding sequences (CDS) from placental tissue MS/MS^65,66^ and the full human UniProt database.

These structural features are consistent with patterns observed in evolutionarily younger genes across multiple species, where gene length and transcript complexity show directional trends toward increasing structural elaboration over evolutionary time.^54^ Approximately half of the novel transcripts in our assembly are non-coding, raising the possibility that some may represent transcriptional noise rather than functional innovations under selection. However, definitive characterization of evolutionary origin and functional significance would require synteny analysis, phylogenetic reconstruction, or experimental validation to distinguish between lineage-specific functional genes, neutrally-evolving transcripts, and transcriptional noise.^55–60^

### The placenta exhibits transcriptional breadth and complexity comparable to other human tissues, challenging its characterization as a transcriptomic void

To evaluate placental transcriptional breadth and complexity with reference to other human tissues, we reprocessed 15 GTEx^47^ tissues and cell lines from raw ONT fastq files using an identical bioinformatic pipeline to our placental samples, including matched assembly, annotation, filtering, and orthogonal validation procedures (**Tables S16-S17**; Methods). Our analysis revealed that the placenta exhibits transcriptional breadth (total number of isoforms and genes expressed) comparable to other human tissues **(Figure 1D)**. Saturation analysis demonstrated that isoform discovery plateaus at similar sequencing depths across all tissues, confirming that our assemblies achieved sufficient depth for robust comparison **(Figure 1E)**. Transcriptional complexity (the mean number of isoforms per gene) ranged from 1.8 to 2.8 across tissues **(Figure 1F)**. These findings challenge previous characterizations of the placenta as a “transcriptomic void”^21,23,61^ expressing few, high-abundance transcripts. Instead, the placenta exhibits transcriptional breadth and complexity comparable to other tissues and utilizes complex splicing regulation—particularly for genes involved in maternal-fetal signaling.

### Pregnancy-related genes exhibit extraordinary transcriptional complexity

While transcriptional breadth and complexity of the placenta are comparable to other tissues, it exhibits exceptional splicing diversity in genes with critical functions in pregnancy and placental development. Chorionic somatomammotropin hormone 1 (*CSH1*), which regulates human placental lactogen production in pregnancy,^62^ displayed the greatest transcriptional diversity with 108 distinct isoforms (**Figure 1F**, maximum values shown in brackets; **Figure 2A-B**). This was followed by pregnancy-specific glycoproteins *PSG5* (56 isoforms) and *PSG6* (48 isoforms), and the CSH-like gene *CSHL1* (50 isoforms). Similarly, growth hormone 2 (*GH2*), which encodes placental growth hormone, also showed expanded transcriptional diversity with 22 distinct isoforms. Both *CSH1* and *GH2* belong to the growth hormone/chorionic somatomammotropin gene family clustered on chromosome 17, and their protein products play coordinated roles in maternal metabolic adaptation to pregnancy and fetal growth regulation. The extensive transcriptional diversity across this gene family suggests complex regulation of placental endocrine functions, supported by significant enrichment for “hormone receptor binding” (odds ratio = 79.3, adjusted p = 1.29 × 10^-3^) for the 50 most isoform-rich genes through Gene Ontology (GO) analysis **(Table S5)**.

### Novel isoforms show varied coding potential and encode potentially functional proteins

We evaluated the coding potential and peptide sequence validation of detected transcripts. Open reading frames (ORFs) were predicted for 19,670 (52.2%) isoforms **(Figure S5A)**, with similar coding sequence (CDS) length across structural categories (median = 1.11 kb) **(Figure S5B)**. Coding potential, measured by CPC2 coding probability scores,^63^ varied by structural category: FSM, NIC and NNC transcripts showed similarly high coding potential (median = 85.39%) while ISM transcripts (median = 66.02%) and other structural categories (median = 57.19%) displayed lower coding potential **(Figure S5C)**.

We used BlastP^64^ to identify isoform-unique peptides from tandem mass spectrometry (MS/MS) of human placental tissue, including GDM-affected pregnancies,^65,66^ ensuring validation of isoform-specific features. We found that 15.2% of reference transcripts (FSM) and 5.0-5.2% of novel isoforms (NIC, NNC) contained unique peptides detected in placental MS/MS **(Figure 2C; Table S6)**. Notably, ISM (22.7%) and genic genomic transcripts (20.0%) showed higher validation rates in placental MS/MS compared to the full UniProt database (9.1% and 10.0%, respectively). In contrast, the full UniProt database revealed higher validation rates for reference and novel junction transcripts (FSM: 31.1%, NIC: 15.0%, NNC: 13.6%), with lower validation in the tissue-specific MS/MS reflecting the limited dynamic range and condition-specific nature of proteomic detection.

We identified 80 transcripts of novel genes (17.62%) with predicted ORFs—44 of which show MS/MS validation— encoding functionally relevant proteins including fibronectin (a potential GDM biomarker),^67^ E-cadherin (regulating trophoblastic differentiation),^68^ alpha-enolase (5 distinct isoforms, differentially expressed in GDM),^69^ occludin (placental barrier integrity),^70^ and CCL14 (trophoblast migration and maternal-fetal communication).^71^ Transcripts without peptide validation may include both non-coding RNAs, which can have important regulatory functions, as well as proteins that may be below MS/MS detection limits due to low abundance, post-translational modifications affecting peptide recovery, condition-specific expression not represented in available proteomic datasets, or cell-type-specific expression diluted in bulk tissue samples.

### Placenta transcript annotations significantly reduce quantification uncertainty in short-read datasets

To evaluate the impact of transcript discovery and filtration on short-read transcript quantification, we compared quantifications against our placenta reference transcriptome, GENCODE v45 and their union (GENCODE+). We analyzed Illumina cDNA libraries from villous placental tissue collected in two independent pregnancy follow-up studies: GUSTO^28^ (n = 200) and the Genetics of Glucose regulation in Gestation and Growth (Gen3G^29^; n = 152), both with approximately 20% of samples from GDM-affected pregnancies **(Figure 3A)**. Following quality control assessment of the sequenced libraries, we removed two GUSTO samples due to low read quality **(Table S7)** and two additional GUSTO samples as statistical outliers **(Figure S6A)**.

**Figure 3.**
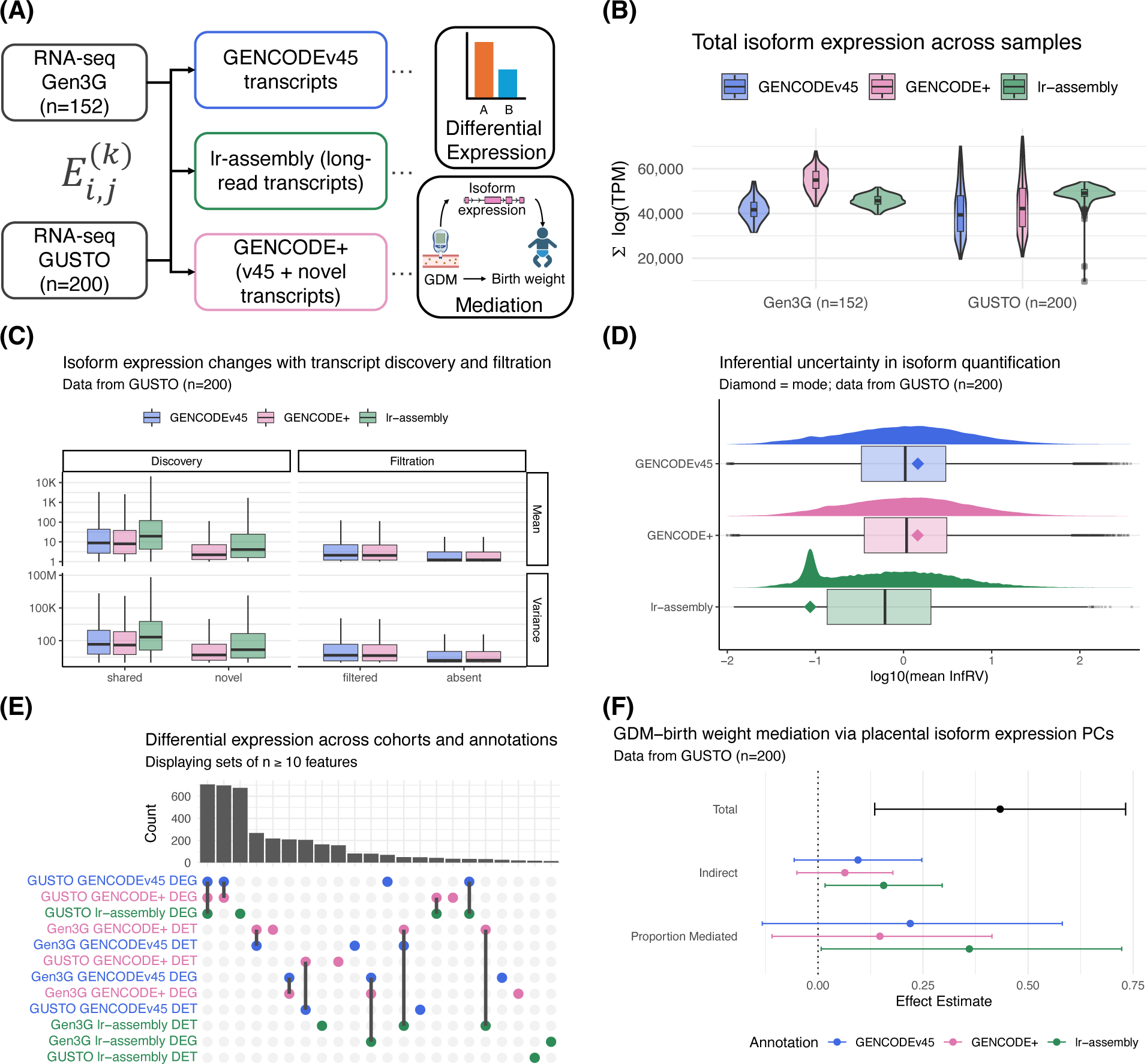
Isoform-aware quantification of placental short-read RNA-seq reduces inferential uncertainty and reveals transcriptional mediation of GDM effects on birth weight. **(A)** Illumina short-read RNA-seq of villous placental tissue from two independent pregnancy follow-up study cohorts with approximately 20% of samples from GDM-affected pregnancies were quantified against the GENCODE v45 transcripts, the high-confidence placenta transcripts (lr-assembly) and their union (GENCODE+). Using Salmon^120^, for each cohort and annotation, the expression *E* of transcript *T_j_* in sample *S_i_* was estimated for *B_k_* bootstrap replicates, where *j* = 1,…,n transcripts in the given annotation, *i* = 1,…,n short-read samples, and *k* = 1,…,50 replicates. Bootstrap replicates were used to estimate overdispersion arising from read-to-transcript mapping ambiguity, which was divided from the counts prior to downstream statistical analyses^25^, and to quantify inferential relative variance (InfRV), a measure of uncertainty in the read-to-transcript mapping^72^. **(B)** The total transcript expression (measured as transcripts per million [TPM]) per short-read sample in two cohorts (GUSTO, n=200; Gen3G, n=152) when quantified against each annotation. **(C)** Comparing mean and variance in the GUSTO short-read samples across references for: transcripts that were shared between annotations (shared), transcripts that were novel in placenta (novel), constituting transcripts included in the placenta reference transcriptome (lr-assembly) through the discovery phase, in addition to GENCODEv45 transcripts that were initially recovered in the long-reads but removed in the filtering procedure (filtered) or that were not assembled from the long-reads (absent), constituting transcripts that were omitted from lr-assembly through the filtration phase. **(D)** Inferential uncertainty (InfRV) in isoform quantification in the GUSTO short-read samples when quantified against transcripts in each annotation. Diamond = mode. **(E)** The count of differentially expressed genes (DEGs) and transcripts (DETs) in each cohort given short-read quantification against each annotation. **(F)** Mediation analysis effect estimates showing the relationship between gestational diabetes mellitus (GDM) and birth weight through placental isoform expression in GUSTO. Principal components were derived from differentially expressed transcripts for each annotation. Mediators included the first *PC* principal components that cumulatively explain at least 75% of expression variance (lr-assembly, PC = 5 [77.09% total variance]; GENCODEv45, PC = 8 [76.72%]; GENCODE+, PC = 6 [75.00%]).

Our placenta reference transcriptome demonstrated population-specific differences in total quantified expression that varied by cohort ancestry. In the GUSTO cohort (Singaporean), which shared ancestry with the samples used to generate the placenta reference transcriptome, lr-assembly achieved the highest total quantified expression with an increase of 7,351 log(TPM) over GENCODE v45 (17.9% increase, p < 2 × 10^-16^) and 4,622 log(TPM) over GENCODE+ (10.6%, p < 2 × 10^-16^) **(Table S8)**. In contrast, the Gen3G cohort (white European) showed the highest total quantified expression with GENCODE+, which combines both annotation sources, with an increase of 13,044 log(TPM) over GENCODE v45 (31.2%, p < 2 × 10^-16^). The lr-assembly alone provided a smaller but significant 3,896 log(TPM) increase over GENCODE v45 in Gen3G (9.3%, p = 1.45 × 10^-10^). These differences in total quantified expression across cohorts (interaction p < 2 × 10-16) likely reflect the ancestry composition of our long-read samples, with GENCODE+ detecting additional isoforms in Gen3G that were not captured in the Singaporean-derived lr-assembly **(Figure 3B)**.

Despite these ancestry-specific differences in total quantified expression, transcripts that failed to meet our stringent quality criteria showed significantly lower mean expression and variance in the short-read quantifications compared to FSM and novel transcripts in both cohorts **(Figure 3C; Figure S6B)**. Statistical analysis revealed a clear hierarchy in transcript expression levels and variance **(Table S9)**: shared (FSM) transcripts showed the highest expression, followed by novel transcripts, then filtered transcripts, and finally absent transcripts. In GUSTO, shared transcripts had 1.14-1.67 log-fold higher expression than other categories (all p < 1 × 10^-4^), with expression variance showing similar patterns (1.88-2.78 log-fold differences, all p < 1 × 10^-4^). Importantly, novel transcripts showed expression levels only slightly lower than shared transcripts, both of which had higher expression levels than filtered or absent transcripts, confirming that our assembly captures functionally relevant novel isoforms rather than low-quality artifacts. Notably, this pattern held in both cohorts: transcripts absent from our lr-assembly but present in GENCODE (which contributed to Gen3G’s higher expression with GENCODE+) also showed significantly lower expression and variance, suggesting many represent low-confidence annotations.

To formally quantify read-to-transcript mapping ambiguity, for each sample across both cohorts, we calculated transcript-level inferential relative variance (InfRV). As InfRV is roughly stabilized with respect to the mean expression level, higher InfRV values indicate greater uncertainty in estimating transcript abundance^72^ (Methods). Mean transcript-level InfRV was consistently reduced when short reads were quantified against our placenta reference transcriptome compared to both GENCODE v45 (30.4% reduced in GUSTO and 30.3% reduced in the Gen3G) and GENCODE+ (31.5% reduced in GUSTO and 30.4% reduced in Gen3G) **(Figure 3D; Figure S6C)**, indicating improved isoform-level quantification precision. Critically, this improvement in quantification precision was observed in both cohorts despite the ancestry-specific differences in total expression, indicating that the lr-assembly reduces mapping ambiguity even for the Gen3G cohort where GENCODE+ achieved higher total quantified expression. When stratified by structural category, transcripts in “Other” structural categories (antisense, intergenic, fusion, and genic genomic) displayed low mean InfRV and the weakest between-sample correlations **(Figure S6D-G).** This pattern suggests unambiguous read mapping due to simpler transcript structures with less exon sharing, but high biological or technical variance reflecting potentially stochastic transcription, sensitivity to cellular state, or lower abundance that amplifies technical noise.

As reduced InfRV is positively correlated with reduced exon sharing between transcripts (**Figure S6H**; r = 0.687, p = 3.85 x 10^-279^), smaller annotations would be expected to show lower InfRV values. To confirm our observed reduction reflected tissue-matched isoform content rather than annotation size alone, we quantified placental short-read RNA-seq against long-read transcripts from two GTEx tissues: subcutaneous adipose (the smallest GTEx assembly with 13,829 transcripts) and cultured fibroblasts (the largest GTEx assembly with 47,850 transcripts). Both GTEx annotations reduced InfRV relative to GENCODE v45 or GENCODE+, consistent with reduced exon sharing from smaller annotations. However, the placenta reference transcriptome yielded substantially lower InfRV values than either GTEx annotation (**Figure S6I**; mode log10(InfRV) estimated by kernel density: placenta = -1.05, adipose = -0.86, fibroblasts = -0.86). This demonstrates that GTEx assemblies contain tissue-specific isoforms not expressed in placenta, introducing irrelevant exon-sharing complexity, while our placenta annotation both excludes irrelevant transcripts through assembly and filtering and includes relevant placental isoforms through discovery. These data confirm that tissue-matched long-read annotations improve quantification precision beyond what would be expected from annotation size reduction alone and that the benefits of our placenta reference transcriptome extend across ancestry groups, even when not all ancestry-specific isoforms are captured.

### Placenta transcript annotations reveal more extensive GDM-associated transcriptional changes

We next examined transcriptional differences at gene and transcript levels associated with GDM status in GUSTO and Gen3G, comparing analyses using our placenta reference transcriptome versus GENCODE v45 and their union (GENCODE+), adjusting for gestational age, fetal sex, and technical variation estimated with RUVr^73^ **(Figure 3E; Figure S7A-B).** Using FDR < 10% thresholds, our placenta reference transcriptome revealed an average of 138 differentially expressed transcripts (DETs) between cohorts (SD = 159.8) and 792.5 differentially expressed genes (DEGs; SD = 955.3), while GENCODE v45 yielded 333.5 DETs (SD = 99.7) and 934.5 DEGs (SD = 839.3) and GENCODE+ yielded 471 DETs (SD = 138.9) and 923 DEGs (SD = 823.1). Analyses using our placenta reference transcriptome resulted in identification of fewer, but a higher annotated percentage of, GDM-associated features **(Tables S10-11)**. This enrichment, combined with the reduced inferential uncertainty we observed across transcripts, suggests that our tissue-specific annotation approach concentrates the analysis on biologically relevant transcripts while filtering out noise from irrelevant or poorly annotated features.

### Placental transcription mediates significant portion of GDM effects on birth weight

We performed pathway enrichment analysis on differentially expressed transcripts (FDR < 10%) in GUSTO, recognizing that different annotation approaches may capture distinct but potentially valid biological signals (Table S12). Our placenta reference transcriptome identified pathways enriched for endocrine and signaling functions: “hormone receptor binding” (odds ratio = 101.89, adjusted p = 3.20 × 10^-3^), “growth factor activity” (odds ratio = 48.42, adjusted p = 2.49 × 10^-3^), and “receptor ligand activity” (odds ratio = 18.81, adjusted p = 2.49 × 10^-3^) **(Figure S7C)**. In contrast, GENCODE annotations revealed pathways related to structural processes: “PI3K-Akt signaling pathway” (odds ratio = 8.19, adjusted p = 7.42 × 10^-7^) and “focal adhesion” (odds ratio = 10.16, adjusted p = 4.23 × 10^-6^) **(Figure S7D)**. To determine which approach better captures functionally relevant changes that mechanistically link GDM to birth weight outcomes, we performed causal mediation analyses.

Given the known impacts of GDM on birth weight, we examined whether some of these effects are mediated by placental transcription in GUSTO using multi-mediator analysis. To address the high-dimensional nature of transcript expression data while maintaining statistical power, we performed principal component analysis on differentially expressed transcripts (FDR < 10%) and, for equal comparison, selected the first *PC* principal components which cumulatively explained at least 75% of the total variance in transcript expression as mediators^74,75^ (lr-assembly, *PC* = 5 [77.09% total variance]; GENCODEv45, *PC* = 8 [76.72%]; GENCODE+, *PC* = 6 [75.00%]). We addressed GDM-transcriptome (exposure-mediator) and transcriptome-birthweight (mediator-outcome) confounding by adjusting for maternal ethnicity, fetal sex, and gestational age. We confirmed these principal components met our 75% variance threshold through sensitivity analysis **(Figure S8A-B; Table S13).** In GUSTO, GDM was significantly associated with increased birth weight (total effect = 0.430, 95% CI [0.131, 0.729], p = 0.005). Importantly, the indirect effect of GDM on birth weight through placental transcription when quantifying short-reads against our placenta reference transcriptome was statistically significant (0.155 [0.016, 0.293], p = 0.029), with approximately 35.8% of the total effect being mediated through placental isoform expression **(Figure 3F)**. Consistent with the pathway enrichment results, our tissue-specific approach identified mediators with clear placental endocrine functions (*CSH1*, *CDC25C*), aligning with the hormone and growth factor pathways detected in the enrichment analysis. In contrast, when analyzing the same samples using either GENCODE v45 or GENCODE+, the proportion mediated by placental transcript expression was smaller and not statistically significant (GENCODEv45: 0.095 [-0.057, 0.247], p = 0.219) (GENCODE+: 0.064 [-0.050, 0.178], p = 0.273) and these annotations produced more heterogeneous mediators including several transcripts of uncharacterized genes.

To confirm that these observations were not merely driven by differences in annotation size, we quantified the GUSTO short-read samples against GTEx transcript assemblies for subcutaneous adipose tissue and cultured fibroblast cells, performing identical differential transcript expression **(Figure S8C)** and multimediator analyses **(Figure S8D-E; Table S13)** as for the other annotations. Consistent with our observation that annotation size alone does not drive differences in quantification uncertainty, quantification against annotations of different sizes yielded insignificant GDM-birth weight mediation by isoform expression (subcutaneous adipose: *PC* = 1, 0.045 [-0.017, 0.106], p = 0.154) (fibroblasts: *PC* = 2, 0.049 [-0.031, 0.128], p = 0.230). These findings demonstrate that both transcript discovery and filtration improve detection of biologically relevant pathway enrichments and estimation of mechanistically relevant transcriptional changes that mediate clinically important outcomes.

### Novel CSH1 isoforms emerge as key mediators of GDM effects on fetal growth

We further identified specific gene isoforms that mediate the effect of GDM on birth weight **(Figure 4A; Table S14)**. In GUSTO, 11 transcripts showed significant mediation effects (p < 0.05), including *ESPRESSO:chr17:12985:70* (0.021, 95% CI [0.001, 0.048], p < 2 × 10^-16^), a novel isoform of *CSH1* not annotated in GENCODE, accounting for approximately 13.4% of the effect of placental transcription. Similarly, in Gen3G, 19 transcripts showed a significant effect, including *ESPRESSO:chr17:12985:62* (0.216 [0.071, 0.389], p < 2 × 10^-16^), a different novel isoform of *CSH1*, accounting for approximately 24.37% of the effect of placental transcription (total effect of GDM on birth weight = 0.132, indirect effect of placental transcription = 0.117). The identification of different *CSH1* isoforms mediating GDM effects across two independent cohorts illustrates the plasticity of alternative splicing in response to genetic and environmental factors **(Figure 4B)**. This population-specific variation was unique to *CSH1* in our placenta reference transcriptome: GENCODE-annotated transcripts showed no significant cross-cohort differences in expression patterns.

**Figure 4.**
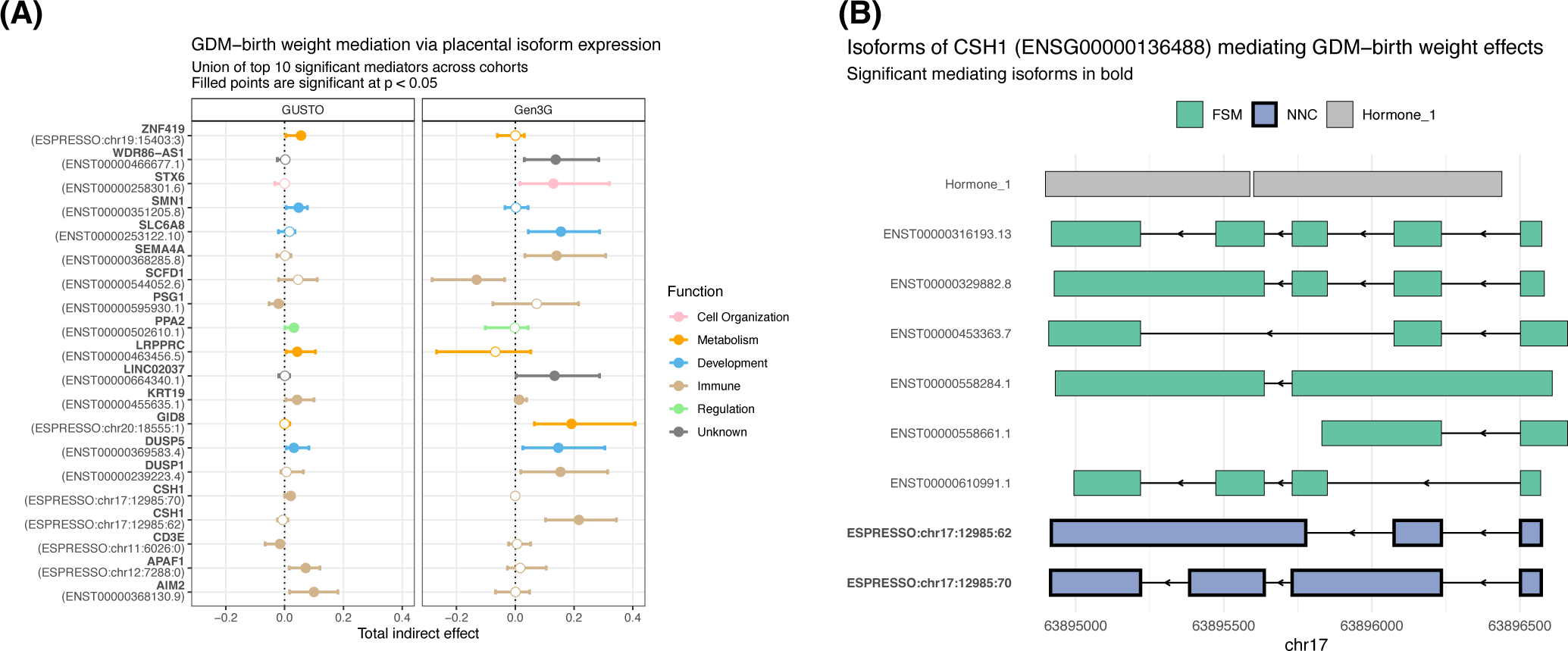
GDM-induced shifts in placental isoform expression are associated with variation in birth weight. **(A)** Individual transcripts showing significant (p < 0.05) GDM-birth weight mediation effects in each pregnancy follow-up study cohort when short-read samples were quantified against the placenta transcripts. **(B)** Transcript structures of the NNC isoforms of *CSH1* showing significant GDM-birth weight mediation effects with reference to the FSM isoforms, illustrating novel intron retention events. Protein domain track for Hormone 1 is indicated in grey.

### Placenta transcript annotations redistribute short-read assignments to dominant and novel isoforms

To understand how differences in transcript annotations affect differential expression (and by extension, causal mediation) analyses, we tracked changes in effect sizes in GUSTO between quantifications on GENCODE v45 and our placenta reference transcriptome **(Figure 5A)**. For *CSH1*, which exhibited the most dramatic expansion in isoform number in our reference, one FSM isoform (ENST00000558284.1) only showed significant differential expression associated with GDM when quantified against GENCODE **(Figure 5B)**. While this isoform showed moderate expression in both references, it had notably higher expression when quantified against GENCODE **(Figure 5C)**, owing to a redistribution of reads to ENST00000558284.1 or to novel isoforms when the data is quantified with our reference **(Figure 5D)**. In contrast, *GNAS*—a known imprinted gene in placenta^76^ that exhibited one of the most dramatic contractions in transcriptional complexity relative to GENCODE (77 isoforms reduced to 17 in our reference)—showed the opposite pattern. Isoform ENST00000476196.5 showed a significant GDM association when quantified against our placenta reference transcriptome but not against GENCODE **(Figure 5E)**, and increased expression with our reference **(Figure 5F-G)**. These findings illustrate how both adding unannotated transcripts and removing irrelevant transcripts from a transcriptome reference results in the redistribution of reads among isoforms of a gene and to other genes. This redistribution is likely beneficial, as it reduces noise by reassigning reads from spurious annotations, improving both quantification accuracy and the detection of biologically relevant differential expression signals.

**Figure 5.**
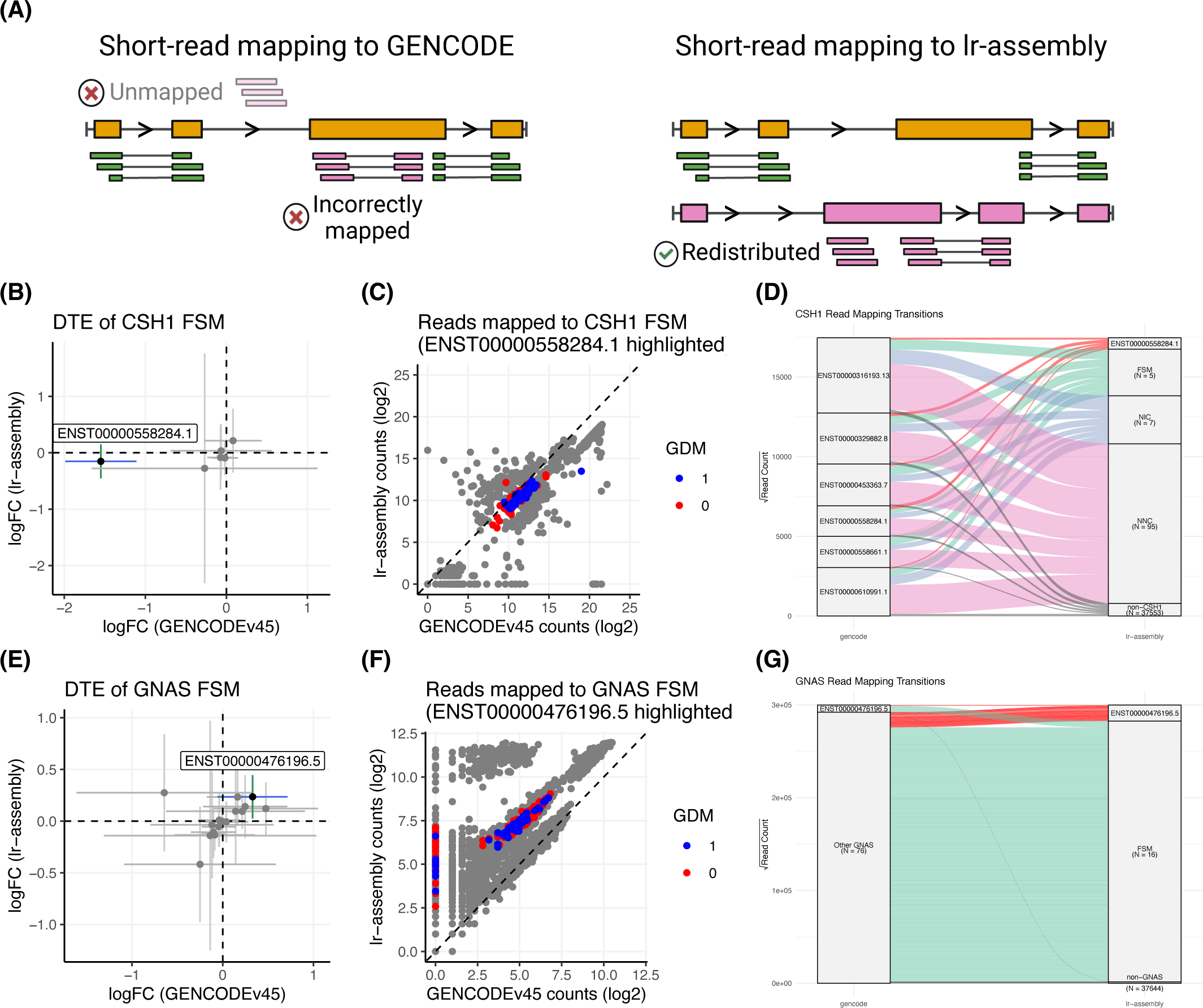
Transcriptome annotation of placenta refines short-read mapping to individual transcript isoforms and uncovers distinct patterns of differential expression. **(A)** Illustration of redistribution of short reads (green and pink boxes) unmapped or incorrectly mapped against GENCODE transcripts (gold transcript) to the placenta reference transcriptome (lr-assembly, gold and pink transcripts). **(B)** Differential transcript expression (DTE) of *CSH1* FSM transcripts in GUSTO. Vertical (green) lines indicate the 95% CI of the log_2_ fold change when quantified against GENCODE v45 transcripts, while horizontal (blue) lines indicate the 95% CI when quantified against lr-assembly transcripts. **(C)** Raw read counts for each *CSH1* FSM transcript for each sample in GUSTO. Highlighted points correspond to ENST00000558284.1, with GDM status indicated. **(D)** Reads mapped to *CSH1* isoforms in GENCODE v45 and their respective mapping locations on the placenta transcript annotations. **(E)** DTE of *GNAS* FSM transcripts in GUSTO. **(F)** Raw read counts for each *GNAS* FSM transcript for each sample in GUSTO. Highlighted points correspond to ENST00000476196.5, with GDM status indicated. **(G)** Reads mapped to *GNAS* isoforms in GENCODE v45 and their respective mapping locations on the placenta transcript annotations.

## Discussion

Despite the placenta’s critical role in fetal programming and later-life health outcomes,^1–7^ it has been notably absent from large-scale genomic initiatives.^18–20^ Using matched bioinformatic processing of placental and GTEx^47^ tissues, our results demonstrate that the placenta exhibits transcriptional breadth and complexity comparable to other adult tissues while displaying exceptional isoform diversity for specific pregnancy-related genes. This pattern challenges previous characterizations of the placenta as a “transcriptomic void”^21,23,61^ and demonstrates that the placenta utilizes complex splicing regulation, particularly for genes involved in maternal-fetal signaling. Gong *et al*. concluded that the placental transcriptome was skewed toward a small number of highly abundant transcripts, suggesting this reflected the placenta’s nature as a rapidly growing and transient organ where many transcripts are “dispensable.” However, our long-read assembly reveals that this apparent simplicity likely reflects the technical limitations of short-read sequencing in resolving complex alternative splicing patterns rather than true biological simplicity. The substantially higher transcriptional complexity we observe may be facilitated by the placenta’s unique DNA methylation landscape.^78–80^ Importantly, transcriptional complexity and breadth metrics are sensitive to sequencing depth, with greater depth yielding more isoforms and genes until saturation is reached. This relationship underscores the importance of adequate sequencing depth and matched processing pipelines for robust cross-tissue comparisons, as depth disparities between studies may lead to spurious differences in apparent transcriptional complexity.

Current placental studies involving transcriptome evaluations rely on gene-level expression measurements, obscuring critical isoform-level regulatory mechanisms and missing key placental functions. This gene-level approach is particularly problematic when adverse outcomes arising from pathological exposures alter functionally distinct gene isoforms—effects that could be masked by compensatory transcriptional responses maintaining total gene expression levels. Critically, our results demonstrate that the choice of reference annotation fundamentally alters differential expression outcomes, with tissue-specific placental annotations revealing biologically meaningful associations that remain hidden when using generic adult tissue references.^77^ Our findings provide empirical support for the benefits of long-read RNA-seq-based isoform-resolved placental analysis by identifying specific placental transcripts, particularly novel *CSH1* isoforms, that mediate the relationship between maternal gestational diabetes mellitus (GDM) and newborn birth weight. These effects would be undetectable in conventional gene-level analyses. By demonstrating that tissue-specific annotations, achieved through both novel transcript discovery and removal of low-confidence transcripts, substantially reduce inferential uncertainty while increasing the interpretability and biological relevance of transcriptomic associations, our work establishes annotation choice as a critical determinant of scientific conclusions in placental genomics and, likely, other tissues.

Our placenta reference transcriptome enables higher-resolution detection of GDM-associated transcriptomic changes. Key placental hormone regulators (*CSH1*, *CSH1L*, *PSG5*, *PSG6*, *GH2*) exhibit remarkable transcriptional complexity, providing flexibility in maternal-fetal hormone signaling. By reducing inferential uncertainty in RNA-seq data,^25,26^ we revealed GDM-linked regulatory changes missed by conventional gene-level approaches.^35–41^ This isoform-resolved precision is critical for identifying early biomarkers of maternal-fetal maladaptation and designing interventions, underscoring the importance of tissue-specific transcriptomic references in developmental contexts.^20–22^ Pathway enrichment analyses using our placenta reference transcriptome revealed GDM-associated transcripts enriched for receptor-ligand activity, hormone receptor binding and growth factor signaling, consistent with key endocrine and paracrine roles of the placenta in orchestrating maternal metabolic adaptations and fetal growth trajectories.^8,9^

Notably, our approach revealed 35.8% of the GDM-birth weight effect was mediated through placental transcription, with population-specific variation in novel *CSH1* isoforms mediating GDM effects across cohorts. *CSH1* encodes human placental lactogen (hPL), a key endocrine regulator of maternal insulin resistance and nutrient partitioning.^10^ The novel *CSH1* isoforms we identified contain intron retention events **(Figure 4B)**, a form of alternative splicing increasingly recognized as an important regulatory mechanism of tissue-specific gene expression,^81,82^ particularly during cellular differentiation^83^ and in response to cellular stress^84^ including aging-induced transposable element expression.^85,86,87^

These findings reinforce the DOHaD hypothesis by identifying isoform-level mediators of maternal-fetal metabolic coupling that reveal population-specific molecular mechanisms linking environmental exposures to gene regulation. Additionally, they provide a potential resolution to previous findings that circulating hPL levels do not correlate with insulin sensitivity changes during pregnancy: studies which measured aggregate hormone levels without accounting for functionally distinct isoforms that may have differential metabolic effects.^42,88,89^ We note that causal mediation analysis assumes no unmeasured exposure-mediator or mediator-outcome confounding, conditional on measured covariates. While this assumption cannot be empirically verified in our study, we minimized potential confounding by adjusting for known confounders of either relationship: maternal ethnicity, fetal sex, and gestational age. Future applications could employ molecular negative control approaches, such as using maternal epigenomic data as proxies for unmeasured confounders,^90^ though such methods require additional structural assumptions.

Our long-read cohort (N = 72) of term placenta represents the largest placental long-read RNA sequencing dataset to date and successfully enabled transcript discovery across two independent validation cohorts with different population ancestries (East/Southeast Asian and White). However, several limitations warrant consideration. The composition of our long-read samples was of Singaporean ancestry, which may limit the capture of ancestry-specific isoforms expressed in other populations. Expanding long-read sequencing to encompass diverse population ancestries will be essential to comprehensively capture population-specific isoform variation and ensure equitable representation in reference transcriptomes. Similarly, future work should expand upon this reference by incorporating placental tissue across stages of gestation and by drawing from larger, more diverse study cohorts with adequate sequencing depth which will be essential for capturing the dynamic regulation of placental transcription throughout development.

A key limitation of bulk tissue analysis is that placental tissue comprises multiple cell types (trophoblasts, endothelial cells, stromal cells, and immune cells), potentially obscuring cell-type-specific isoform expression. The isoform diversity we observe likely represents aggregate expression across these cell types, and cell-type-specific isoform switching or differential isoform usage cannot be resolved in our current dataset. Additionally, while long-read sequencing excels at transcript structure discovery, it has lower throughput than short-read sequencing, which we addressed by using our long-read-defined transcriptome as a reference for isoform-level quantification in larger cohorts with short-read sequencing data. Expanding long-read sequencing to additional populations, trimesters, and cell types, including single-cell or spatial transcriptomics approaches that can resolve cell-type-specific isoform expression, will reveal additional population-, pregnancy stage-, and cell-type-specific isoforms not captured in our current reference.

Experimental validation is also necessary to confirm that the novel protein-coding genes identified here are actively translated and to determine their contributions to placental function. Furthermore, functional follow-up studies of genes like *CSH1* require precise isoform-level resolution, as experimental interventions like transcriptional knockdown or overexpression depend on targeting the exact isoform mediating the biological effect. Finally, while our long-read sequencing approach provides high-confidence transcript structures, future validation studies could employ PCR-free methods such as Oxford Nanopore direct RNA sequencing or spatial transcriptomics approaches to provide independent technical confirmation and tissue localization of novel isoforms.

We also acknowledge that RNA integrity in term placental samples collected under clinical conditions is relatively but expectedly lower than standard quality thresholds. Placental samples must be assessed by a pathologist before collection and transport to the core facility or laboratory handling the RNA extractions, a process that can last 3-4 hours and result in RIN values below the typical threshold for library preparation.^91^ While our samples showed moderate RNA degradation (long-read samples, mean RIN = 6.95 ± 0.77; GUSTO short-read samples, mean = 3.56 ± 1.60; Gen3G short-read samples, mean = 8.3 ± 0.90), RIN values did not differ significantly by GDM status **(Table S15),** ensuring that our differential expression and mediation findings are not confounded by differences in RNA quality. Long-read sequencing is also robust to moderate RNA degradation, and our stringent filtering criteria ensured that only well-supported transcripts were included in downstream analyses.

Our placental transcriptome reference will enable more precise investigation of environmental exposure effects by identifying specific mechanistic pathways that mediate exposure-outcome relationships. By resolving isoform-level mediators, our approach enables identification of less confounded mechanistic pathways that can support causal inference methods, complementing Mendelian randomization approaches by focusing on intermediate biological mechanisms rather than genetic instruments. This mechanistic precision is particularly valuable for studying environmental exposures like maternal diabetes, where effects may operate through distinct gene isoforms with opposing functions. Importantly, our tissue-specific reference reduces the overall transcript annotation set while enriching for biologically relevant isoforms, facilitating fine-mapping of quantitative trait loci (QTLs) and enabling more precise prioritization of causal “effect isoforms” at specific loci for targeted functional validation studies. Furthermore, our demonstration of population-specific transcript mediators establishes a framework for investigating differential environmental exposure effects by ancestry and context, enabling identification of population-tailored intervention targets and biomarkers. Beyond exposure studies, this resource provides a foundation for isoform-resolved association studies of developmental traits and birth outcomes, enabling precise mapping of regulatory variants influencing fetal growth and early-life disease risk.^5,36–38,92,93^ It will enhance comparative evolutionary transcriptomics by improving identification of lineage-specific innovations, such as transposon exonization^94,95^ and transposon-driven alternative promoters,^96,97^ that have shaped placental evolution.^33,86,87,98^ Additionally, it will support detailed investigations of imprinting and parent-of-origin effects,^99^ which are enriched in placental tissues and contribute substantially to pregnancy outcome variation.^76,100–102^ To facilitate exploration of our placental transcriptome resource, we developed an interactive Shiny application (https://github.com/sbresnahan/lr-placenta-transcriptome-viz) that enables users to visualize transcript structures, query gene-level and isoform-level expression patterns, and explore differential expression and mediation analysis results.

## Materials and Methods

### Biological samples and sequencing

We included a subset of two mother-offspring cohort studies^28,52^ of term, live births (N = 72) without known placentation pathologies, such as intrauterine growth restriction and pre-eclampsia, and 50% from pregnancies affected by gestational diabetes mellitus (GDM; categorized by a 75 g two-timepoint oral glucose tolerance test using the WHO 1999 criteria which was in clinical use at that time). Placentae were processed within 1 hr of delivery. Five villous biopsies of each placenta were rinsed with phosphate buffer saline (PBS), snap-frozen with liquid nitrogen and stored at −80 °C before RNA extraction. Of the 72 long-read samples, N = 6 were selected from the Growing Up in Singapore Toward Healthy Outcomes (GUSTO^28,103^) cohort and have matched Illumina short-read libraries from the same biological specimens; the remaining N = 66 long-read samples were selected from an independent National University of Singapore Hospital cohort.^52^ For the N = 6 GUSTO samples, RNA was extracted from approximately 100 mg of crushed tissue using phenol-chloroform and purified with a Qiagen RNeasy Mini Kit. For the N = 66 samples selected from our independent pregnancy follow-up study cohorts, approximately 75 mg of crushed placenta tissue was combined with 500 μL Buffer ATL (Qiagen, #939011) and 30 μL Proteinase K (from Qiagen QIASymphony DSP DNA Midi Kit) and incubated overnight in a shaking incubator. Extracted samples were sequenced via 3x 24-plex on PromethION (release 23.04.6, Oxford Nanopore, ONT) with manufacturer’s PCR-cDNA library prep kit (#SQK-PCB111.24) at the Genome Institute of Singapore using manufacturer’s protocols. Median read lengths were approximately 1,000 bp (vs. 150 bp via NovoSeq).

Paired-end Illumina libraries of villous placenta from term, live births from GUSTO (N = 200) and Genetics of Glucose Regulation in Gestation and Growth^29,104^ (Gen3G, N = 152; phs003151) were sequenced previously. Briefly, placental villous tissue samples were collected after delivery and stored at -80 °C. In total RNA extracted using phenol-chloroform followed by purification with the NucleoSpin RNA kit (Macherey-Nagel). In GUSTO, RNA quality was assessed by Nanodrop spectrophotometry and Agilent TapeStation, with ribosomal RNA depletion performed using Illumina Ribo-Zero kits. Sequencing libraries were prepared using NEB Next Ultra Directional RNA Library Prep Kit and sequenced on Illumina HiSeq 4000 platform to a minimum of 50 M paired-end 150 bp reads per library. In Gen3G, RNA quality was assessed using an Agilent Bioanalyzer and sent to the Broad Institute for sequencing. Libraries were prepared from 250 ng of RNA using Illumina TruSeq Stranded mRNA Sample Preparation Kits and sequenced on an Illumina HiSeq 4000 platform to a minimum of 14 M paired-end 100-bp reads per library.

### Transcriptome assembly

FAST5 files of the raw ONT signals for each sample were processed with Guppy v.6.5.7 for base calling on a GPU with barcode adapter trimming enabled. FASTQ files labeled “pass” were concatenated together for each sample. Pychopper2 was used to identify, orient and trim full-length cDNA reads containing primer sequences used in library construction with the PCS111 sequencing kit (ONT). Full length and rescued cDNA reads were concatenated for each sample, and microindels and sequencing errors were corrected with FMLRC2^105^ using short-read RNA-seq samples of villous placenta from GUSTO (N = 200).^28^ The corrected reads were aligned to the human genome assembly analysis set (GenBank assembly 000001405.15, no ALT contigs, hg38) using Minimap2^106^ with a kmer length of 14, minimizer window size of 4, and in splice-aware mode with --splice-flank=no to relax assumptions about bases adjacent to splice junction donors and acceptors. Secondary alignments were filtered, and primary alignments were coordinate sorted with samtools.^107^ The filtered and sorted alignments to chromosomes 1-22, X, Y and M for all samples were then concatenated together to generate a single BAM file used for de novo transcript assembly with ESPRESSO, which shows favorable balance in maximizing discovery while limiting false discoveries.^49^

### Transcriptome annotation and filtering

The assembled transcript models were filtered using a custom R^108^ script to remove isoforms with un-stranded exons (retaining only “+” or “-” strand isoforms), isoforms with an exon less than 3 nucleotides long, and isoforms with exons positions outside the transcript start and end positions. We then used SQANTI3^109^ to characterize the *de novo* assembly and classify isoforms into structural categories relative to GENCODE v45 transcripts: full splice matches (FSM), incomplete splice matches (ISM), novel in catalog (NIC), novel not in catalog (NNC), and other classes (e.g. fusion products and antisense transcripts). Open reading frames and coding sequences were predicted with TransDecoder.^110^ Coding potential was calculated with CPC2.^63^ For protein validation, we performed BLASTP^64^ searches of predicted peptide sequences against two databases: (1) peptides identified by tandem mass spectrometry (MS/MS) of placental tissue samples, including from GDM-affected pregnancies,^65,66^ and the full UniProt human proteome. To ensure isoform-specific validation, we extracted the matched peptide sequences from each BLASTP hit and identified peptides unique to individual isoforms within each gene. Only transcripts with at least one unique peptide match (> 95% sequence identity) and CPC2 coding probability > 0.5 were considered validated. Tissue-specific gene enrichment was assessed with TissueEnrich.^53^

For quality control, we considered long-read coverage of the assembled transcripts quantified with ESPRESSO, in addition to splice junction coverage in the short-read RNA-seq samples, proximity of transcription start sites (TSS) from Cap Analysis of Gene Expression sequencing (CAGE-seq) peaks, genome-wide sequencing of regions sensitive to cleavage by DNase I (DNase-seq) peaks or reference TSS, and proximity of transcription termination sites (TTS) from sequencing alternative polyadenylation (polyA) sites (SAPAS), polyA motifs or reference TTS **(Table S16)**. We defined high-confidence isoforms across all structural categories as those with a minimum of 3 long (Nanopore) reads covering the transcript and at least 3 short (Illumina) reads covering each splice junction. We validated transcription start sites (TSS) by requiring them to be within 5 kb of a CAGE-seq peak in the FANTOM5 placenta dataset,^111^ within 5 kb of a DNase-seq peak in the ENCODE placenta dataset,^19^ or within 100 bp of a reference TSS. Similarly, transcription termination sites (TTS) were required to be within 100 bp of a SAPAS site in the APASdb placenta dataset,^112^ within 100 bp of a reference TTS, or within 40 bp of a PolyA motif in the PolyA_DB dataset.^113^ For all categories except FSM, we required transcripts to have less than 80% adenosine content in the 20 nucleotides at the 3’ end to eliminate artifacts of PolyA intrapriming.^109^ For novel isoform categories (excluding FSM, ISM and NIC), we applied additional filters: removing transcripts with evidence of reverse transcriptase (RT) switching at splice junctions (a known technical artifact) and applying additional scrutiny to transcripts predicted to undergo nonsense-mediated decay (NMD) including substantial 5’ expression (defined as a ratio of > 0.5:1.0 between short-reads mapping to the first and last splice junctions) to distinguish genuine NMD substrates from assembly artifacts.

### Preparation of orthogonal datasets for assembly filtering

Short-read RNA-seq reads from term, villous placenta from GUSTO (N = 200) were aligned to hg38 using STAR^114^ with settings recommended by ENCODE. Splice junctions with < 10 supporting reads were filtered. CAGE-seq reads from placenta tissue (N = 1) and cells (pericyte, N = 2; epithelial, N = 4, trophoblast, N = 1) were retrieved from the FANTOM5 Project^111^, trimmed with cutadapt v4.1^115^ to remove linker adapters, EcoP15 and 5’poly-G sequences, and filtered reads were aligned to h38 using STAR with settings recommended by ENCODE **(Table S16)**. CAGE tags were counted and clustered using CAGEr^116^ in R. DNAse-seq peaks from placenta tissue (N = 22) mapped to h38 were retrieved in BED format from the ENCODE consortium and intersected with bedtools.^117^ SAPAS peaks in placenta tissue mapped to hg38 were retrieved in BED format from APASdb,^112^ and human polyA motifs annotated in PolyA_DB^113^ were retrieved from PolyA-miner.^118^

### Comparative Analysis of GTEx Tissues with Matched Processing Pipeline

To enable direct comparison of placental transcriptional complexity with other human tissues, we reprocessed 15 GTEx^47^ v9 tissues and cell lines from raw Oxford Nanopore Technologies (ONT) long-read RNA sequencing fastq files (retrieved from dbGaP, phs000424.v10.p2) using an identical bioinformatic pipeline to our placental samples. Assembled transcripts were annotated using SQANTI3 with the same reference files and parameters as placental samples, including GENCODE v45 comprehensive annotation and the human polyA motifs database. For splice junction validation and transcript filtering, we utilized tissue-matched short-read RNA-seq samples from GTEx (retrieved from dbGaP, phs000424.v10.p2) and orthogonal datasets from multiple public repositories (FANTOM5, ENCODE, SRA, PolyA-Site) employing diverse sequencing methods (CAGE, RAMPAGE, 3’-seq, SAPAS, PolyA-seq, DRS) **(Table S16)**. These orthogonal datasets were processed identically to placental validation data. We applied identical SQANTI3 filtering criteria to all tissues to retain high-confidence isoforms. Long-read sample sizes, read depth metrics, and assembly statistics for all GTEx assemblies are provided in **Table S17**.

### Short-read RNA-seq analyses

Paired-end Illumina libraries of villous placenta from term, live births of the GUSTO (N = 200) and Gen3G were sequenced previously. We used fastp^119^ to trim sequencing adapters and filter low-quality reads. Decoy-aware transcriptome indices for Salmon^120^ for GENCODE v45 transcripts, our long-read defined placental transcripts (lr-assembly), and their union (GENCODE+) were constructed with MashMap.^121^ Transcriptome quantification was then performed with Salmon with settings --libType A --validateMappings --seqBias and including 50 bootstrap replicates to estimate:

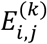

the expression *E* of transcript *T_j_* in sample *S_i_* for *B_k_* bootstrap replicates, where *j* = 1,…,n transcripts in the given assembly, *i* = 1,…,n short-read samples, and *k* = 1,…,50 replicates. Additional quantifications were performed against the long-read transcript assemblies of GTEx^47^ v9 subcutaneous adipose tissue and cultured fibroblast cells following the same procedure described above.

Transcript quantifications were loaded into R with tximport,^122^ and overdispersion arising from read-to-transcript mapping ambiguity was estimated with catchSalmon() from the edgeR package^25^ and divided from the counts^25^ prior to downstream analyses. To reduce noise in downstream analyses, we filtered the expression matrices to retain only transcripts with at least 5 read counts in a minimum of 3 samples. Pearson correlation coefficients were calculated for each pair of samples within each cohort on the transcripts-per-million (TPM) normalized counts.

To test for differences in total transcript expression (TPMs) between assemblies, we used linear mixed-effects models with assembly and cohort as fixed effects and sample as a random effect. To examine differences in mean and variance properties between assemblies and transcript sets, we performed nested ANOVAs using log-transformed mean or variance in expression as the outcome variable, with cohort-by-transcript-set and cohort-by-assembly-set interaction terms. Post-hoc pairwise comparisons between transcript sets within each cohort were conducted using estimated marginal means with Tukey adjustment for multiple comparisons.

Differential transcript and gene expression analyses of GDM status were performed with DESeq2.^123^ Models for all analyses adjusted for gestational age (scaled), fetal sex and 2 dimensions of residual variation estimated with RUVr^73^. Isoform-level mediation analyses^124^ of the GDM-associated differentially expressed transcripts at FDR <10% were performed with the mediation package^125^ using variance stabilizing transformation (VST) normalized counts to estimate the indirect effect of GDM on birth weight (scaled).

Multi-mediator analyses were performed with the lavaan package^126^ selecting as mediators, for equal comparison, the first *PC* principal components^74^ that cumulatively explained at least 75% of the total variance in the VST-normalized counts of the GDM-associated transcripts (DTEs). Models estimated the total effect of GDM and the indirect effect of placental transcription on birth weight, adjusting for gestational age and fetal sex to. To assess potential confounding by population structure, we used one-way ANOVA to test for associations between each PC and maternal ethnicity, calculating p-values and eta-squared effect sizes using the effectsize package^127^ to quantify association strength.

Transcript-level Inferential relative variance (InfRV) across median-ratio scaled bootstrap replicates^72^ was calculated as:

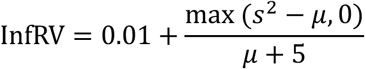

where 𝑠^2^ is the sample variance of scaled counts over bootstrap replicates, and 𝜇 is the mean scaled count over bootstrap replicates, with a pseudocount of 5 added to the denominator and 0.01 added overall to stabilize the statistic and make it strictly positive for log transformation. To visualize read mapping transitions between assemblies for N = 10 randomly selected samples from GUSTO (with 50% from GDM affected pregnancies), we aligned the short-reads with BWA-MEM2^128^ to the transcript sequences in each assembly with settings -O 12,12 -E 4,4 to prevent gapped alignments. End-to-end alignments to the transcript sequences were then filtered with a custom Perl script to retain only the first mate in the read pair and to remove secondary and supplementary alignments.

### Ethics Approval

The Singaporean studies follow guidelines set forth in the Declaration of Helsinki and all study activities were approved by the National Healthcare Group Domain Specific Review Board and the SingHealth Centralized Institutional Review Board (GUSTO only). Informed written consent was obtained from all mothers at recruitment. The ethical committee from the Centre Hospitalier Universitaire de Sherbrooke (CHUS) reviewed and approved Gen3G study protocols.

## Acknowledgements

We thank members of both the Bhattacharya and Huang labs for critical reading of our manuscript, and members of the Love lab for feedback on methods.

## Data Availability

We have made our high-confidence placenta reference transcriptome publicly available at https://github.com/sbresnahan/lr-placenta-transcriptome. An interactive Shiny application for visualizing transcript structures and exploring differential expression and GDM-birth weight mediation results is available at https://github.com/sbresnahan/lr-placenta-transcriptome-viz. Oxford Nanopore reads generated in this study are accessible through the NCBI Sequence Read Archive (BioProject accession PRJNA1373130). Illumina short reads generated through the Gen3G study can be accessed through dbGaP (phs003151.v1.p1), and those generated through the GUSTO study can be accessed with permission from the data access committee (https://gustodatavault.sg/about/request-for-data).

## Code Availability

All custom scripts and analysis pipelines used in this study are publicly available at https://github.com/sbresnahan/lr-placenta-transcriptome. This includes code for dataset preparation, Oxford Nanopore data processing and transcriptome assembly using ESPRESSO, SQANTI3-based annotation and quality control filtering, and short-read analyses in pregnancy follow-up study cohorts including differential gene and transcript expression, mediation analysis, and GO enrichment analysis.

## Author Contributions

S.T.B., J.Y.H and A.B. designed the study. H.Y. and J.K.Y.C. prepared samples for sequencing. M.-F.H. managed the Gen3G study and S.-Y.C. managed the GUSTO study. S.T.B. performed all bioinformatic and statistical analyses and wrote the initial draft of the manuscript. All authors contributed to the writing and editing of the final manuscript.

## Tables

**Table 1.**
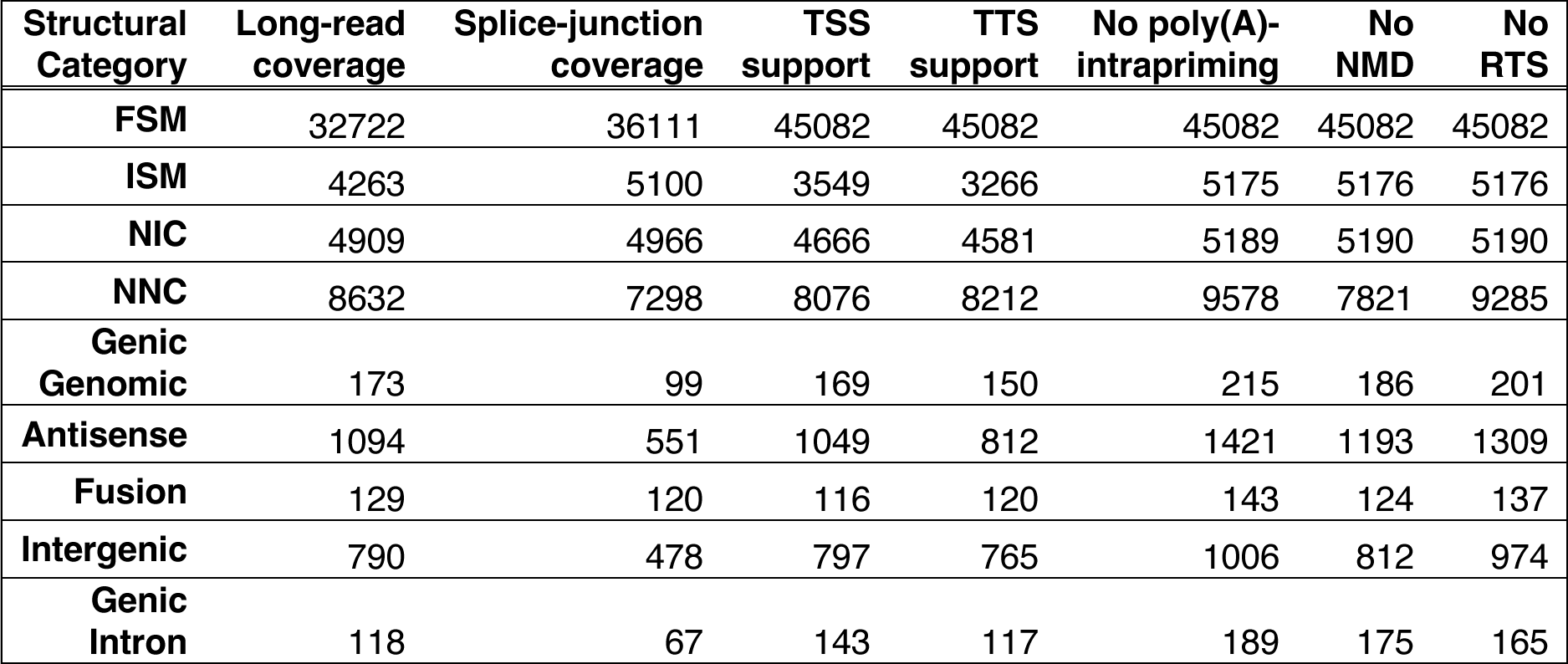
Summary of orthogonal support for placenta transcripts assembled from long-read sequencing data. Stratified by transcript structural category relative to GENCODEv45, the count of isoforms passing thresholds for long-read coverage, short-read coverage of splice junctions, with TSS and TTS support, and without evidence of poly(A)-intrapriming, nonsense mediated decay (NMD) or reverse template switching (RTS) are indicated.

| **Structural Category** | **Long-read coverage** | **Splice-junction coverage** | **TSS support** | **TTS support** | **No poly(A)-intrapriming** | **No NMD** | **No RTS** |
| --- | --- | --- | --- | --- | --- | --- | --- |
| **FSM** | 32722 | 36111 | 45082 | 45082 | 45082 | 45082 | 45082 |
| **ISM** | 4263 | 5100 | 3549 | 3266 | 5175 | 5176 | 5176 |
| **NIC** | 4909 | 4966 | 4666 | 4581 | 5189 | 5190 | 5190 |
| **NNC** | 8632 | 7298 | 8076 | 8212 | 9578 | 7821 | 9285 |
| **Genic Genomic** | 173 | 99 | 169 | 150 | 215 | 186 | 201 |
| **Antisense** | 1094 | 551 | 1049 | 812 | 1421 | 1193 | 1309 |
| **Fusion** | 129 | 120 | 116 | 120 | 143 | 124 | 137 |
| **Intergenic** | 790 | 478 | 797 | 765 | 1006 | 812 | 974 |
| **Genic Intron** | 118 | 67 | 143 | 117 | 189 | 175 | 165 |

