## Supplementary Dataset 2 for "Long-read transcriptome assembly reveals vast transcriptional complexity in the placenta associated with metabolic and endocrine function"

**Supplementary Data 2: Long-read transcriptome assembly reveals vast transcriptional complexity in the placenta associated with metabolic and endocrine function**

Sean T. Bresnahan[1*], Hannah Yong[2], William H. Wu[3], Sierra Lopez[4], Jerry Kok Yen Chan[5,6], Frédérique White[7], Pierre-Étienne Jacques[7], Marie-France Hivert[8,9], Shiao-Yng Chan[2,10], Michael I. Love[11], Jonathan Y. Huang[2,4,6⸸], Arjun Bhattacharya[1,12⸸]

[1]The University of Texas MD Anderson Cancer Center, Department of Epidemiology, Houston, TX
[2]A*STAR Institute for Human Development and Potential, Singapore, Singapore
[3]Rice University, Department of BioSciences, Houston, TX
[4]The University of Hawaiʻi at Mānoa, Department of Public Health Sciences, Honolulu, HI
[5]KK Women's and Children's Hospital, Singapore, Singapore
[6]Duke-NUS Medical School, Singapore, Singapore
[7]Department of Biology, Sherbrooke University, Sherbrooke, QC, Canada
[8]Diabetes Unit, Department of Medicine, Massachusetts General Hospital, Boston, MA
[9]Department of Medicine, Sherbrooke University, Sherbrooke, QC, Canada
[10]National University of Singapore, Yong Loo Lin School of Medicine, Department of Obstetrics & Gynecology, Singapore, Singapore
[11]University of North Carolina–Chapel Hill, Departments of Genetics & Biostatistics, Chapel Hill, NC
[12]The University of Texas MD Anderson Cancer Center, Institute for Data Science in Oncology, Houston, TX

[⸸]Co-last authors

### Contents

Figure S1 – Raw assembly summary statistics
Figure S2 – Quality control of transcript models
Figure S3 – High-confidence assembly summary statistics
Figure S4 – Known vs novel transcript features
Figure S5 – Coding sequence features
Figure S6 – Short-read quantification analyses
Figure S7 – Differential expression analyses
Figure S8 – Isoform expression causal mediation analyses

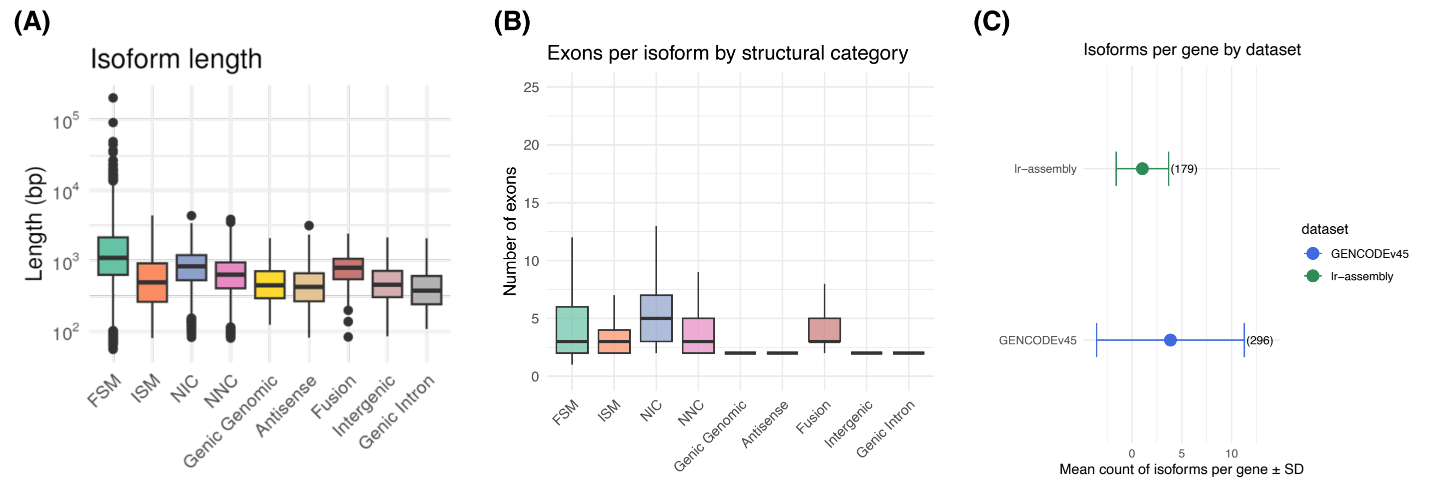


**Figure S1** – **Raw assembly summary statistics**. **(A)** Isoform length distribution by structural category. **(B)** Exons per isoform by structural category. **(C)** Isoforms per gene in GENCODEv45 versus the placenta transcriptome reference. Maximum values shown in brackets.

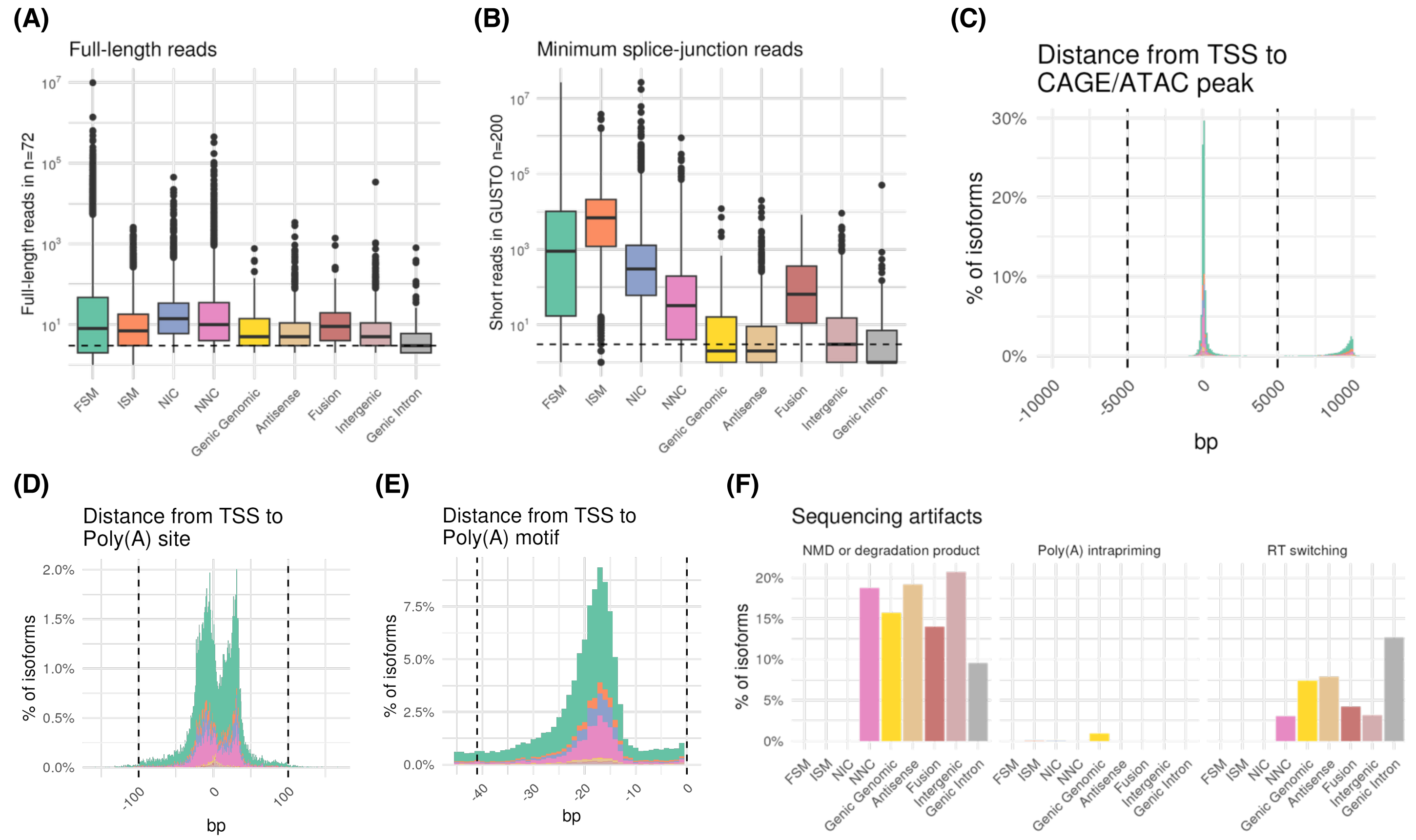


**Figure S2** – **Quality control of transcript models**. **(A)** Long-read support. **(B)** Splice junction support in the GUSTO short-read dataset. **(C)** CAGE/DNase-seq support of TTS. **(D)** SAPAS support of TTS. **(E)** TTS distance from Poly(A) motifs. **(F)** Sequencing artifacts: nonsense-mediated decay (NMD), RT-switching, Poly(A)-intrapriming.

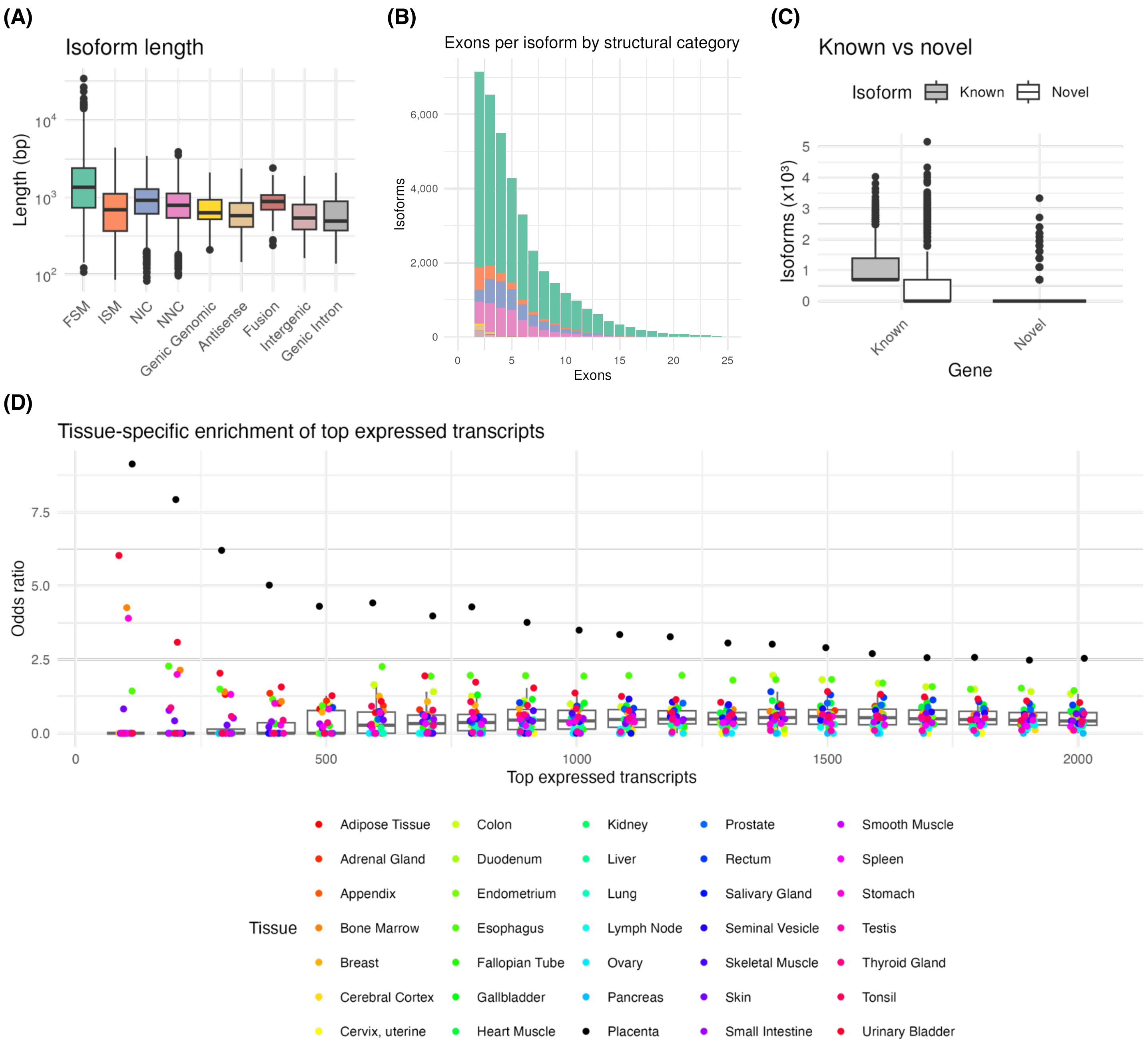


**Figure S3** – **High-confidence assembly summary statistics**. **(A)** Isoform length distribution by structural category. **(B)** Exons per isoform by structural category. **(C)** Genes and isoforms, stratified by known (annotated in GENCODEv45) vs novel in the placenta transcriptome reference. **(D)** Tissue-specific enrichment of top expressed transcripts.

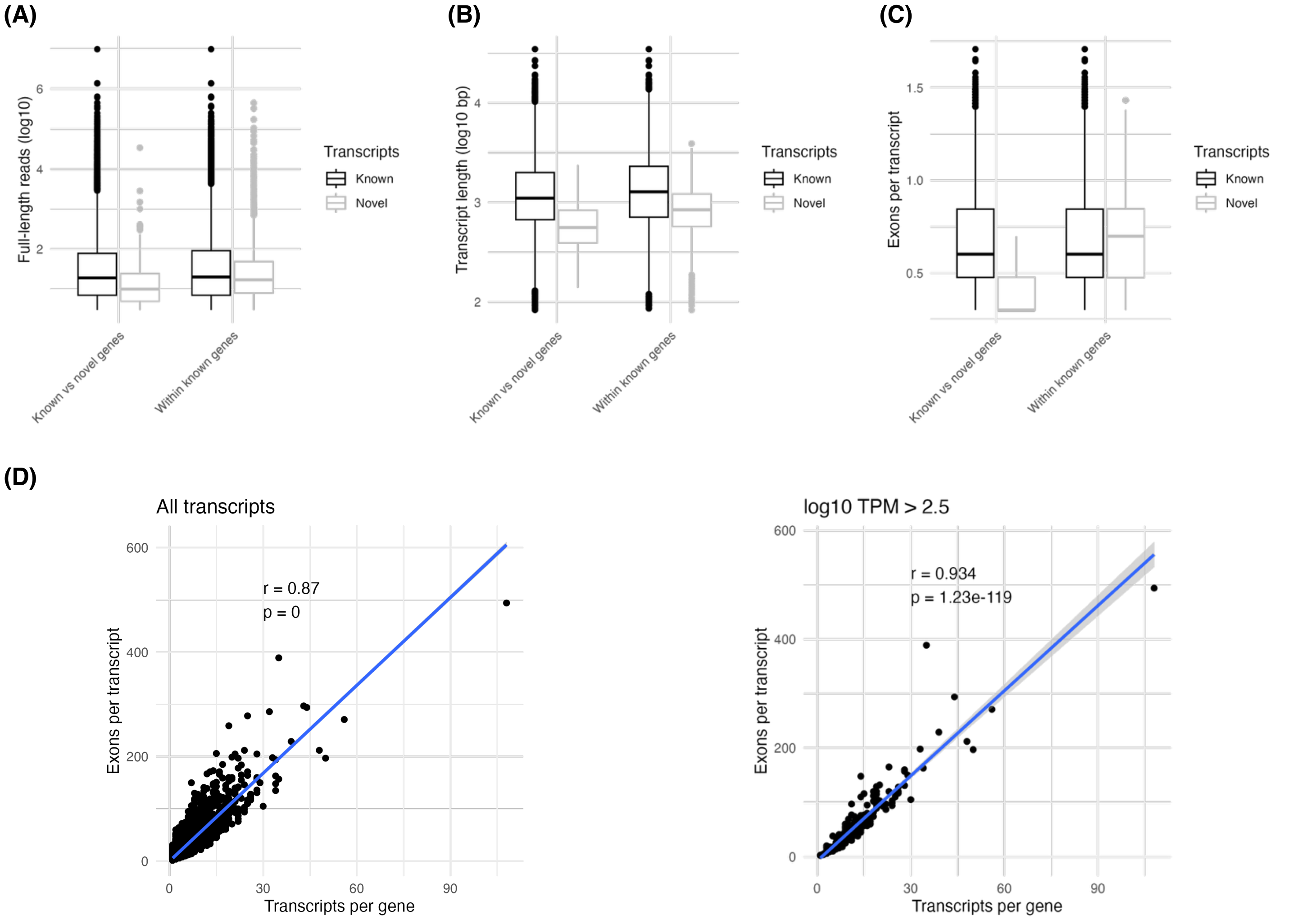


**Figure S4** – **Known vs novel transcript features**. Comparisons between transcripts of known vs novel genes, and between FSM transcripts and novel transcripts of known genes in **(A)** long-read read coverage, **(B)** transcript lengths, and **(C)** number of exons. **(D)** Correlation between number of detected isoforms and number of detected exons per isoform.

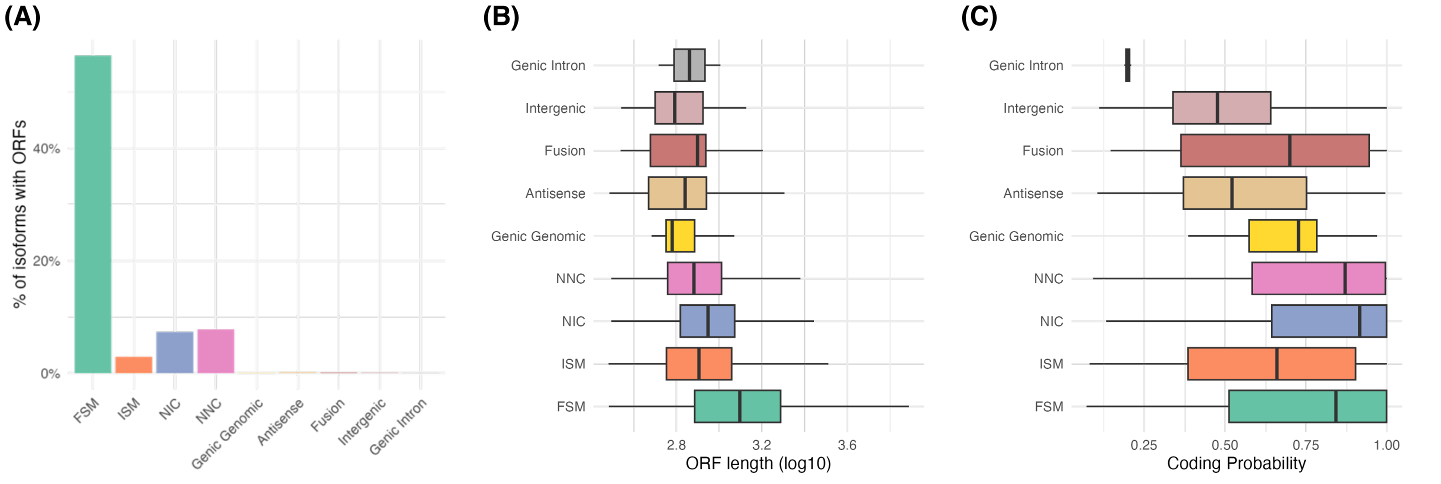


**Figure S5** – **Coding sequence features**. **(A)** Open reading frames (ORFs) predicted per isoform structural category. **(B)** ORF length per isoform structural category. **(C)** CPC2 coding probability scores by isoform structural category.

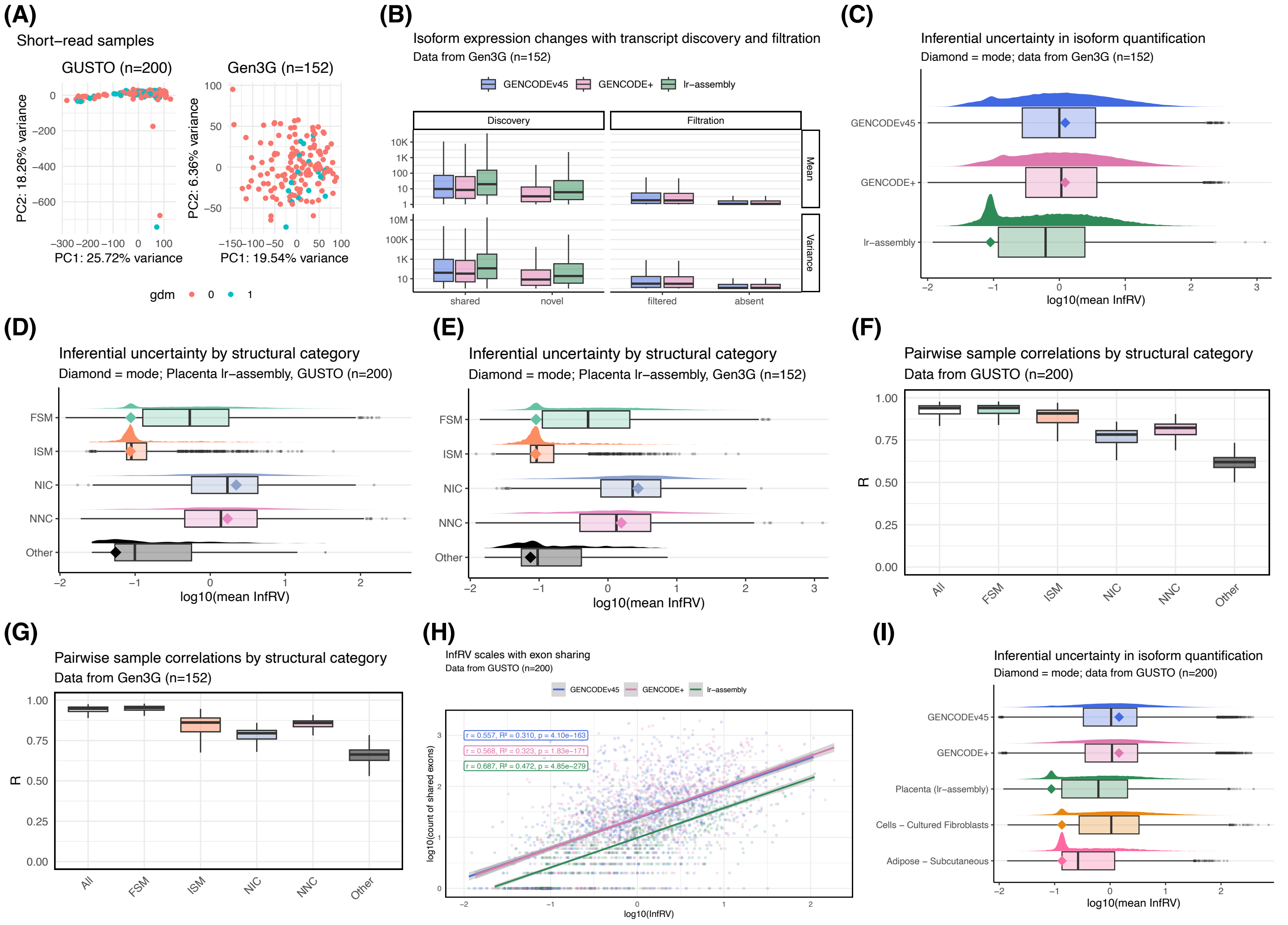


**Figure S6** – **Short-read quantification analyses**. **(A)** Sample PCA of the Variance Stabilizing Transformation normalized counts in each cohort when short-read samples were quantified against the placenta transcriptome reference. **(B)** Isoform expression mean and variance properties in Gen3G when short-read samples were quantified against the placenta transcriptome reference (lr-assembly), GENCODEv45 or their union (GENCODE+). **(C)** Inferential relative variance (InfRV) in transcript quantification of Gen3G short-read samples. **(D)** InfRV by transcript structural category in GUSTO. **(E)** InfRV by transcript structural category in Gen3G. **(F)** Pairwise correlations of transcript expression by isoform structural category in GUSTO. **(G)** Pairwise correlations of transcript expression by isoform structural category in Gen3G. **(H)** Linear relationship between InfRV and exon sharing between transcripts within genes (the number of overlapping exons, calculated for each transcript) in GUSTO. Pearson correlation coefficient (r), $R^2$ and correlation p-value are indicated for short-read quantifications against each annotation. **(I)** In GUSTO, InfRV across all transcripts given short-read quantifications against the placenta transcriptome reference, GENCODEv45 and their union (GENCODE+). Long-read GTEx transcript assemblies for subcutaneous adipose tissue and cultured fibroblast cells provided as references. Diamond = mode.

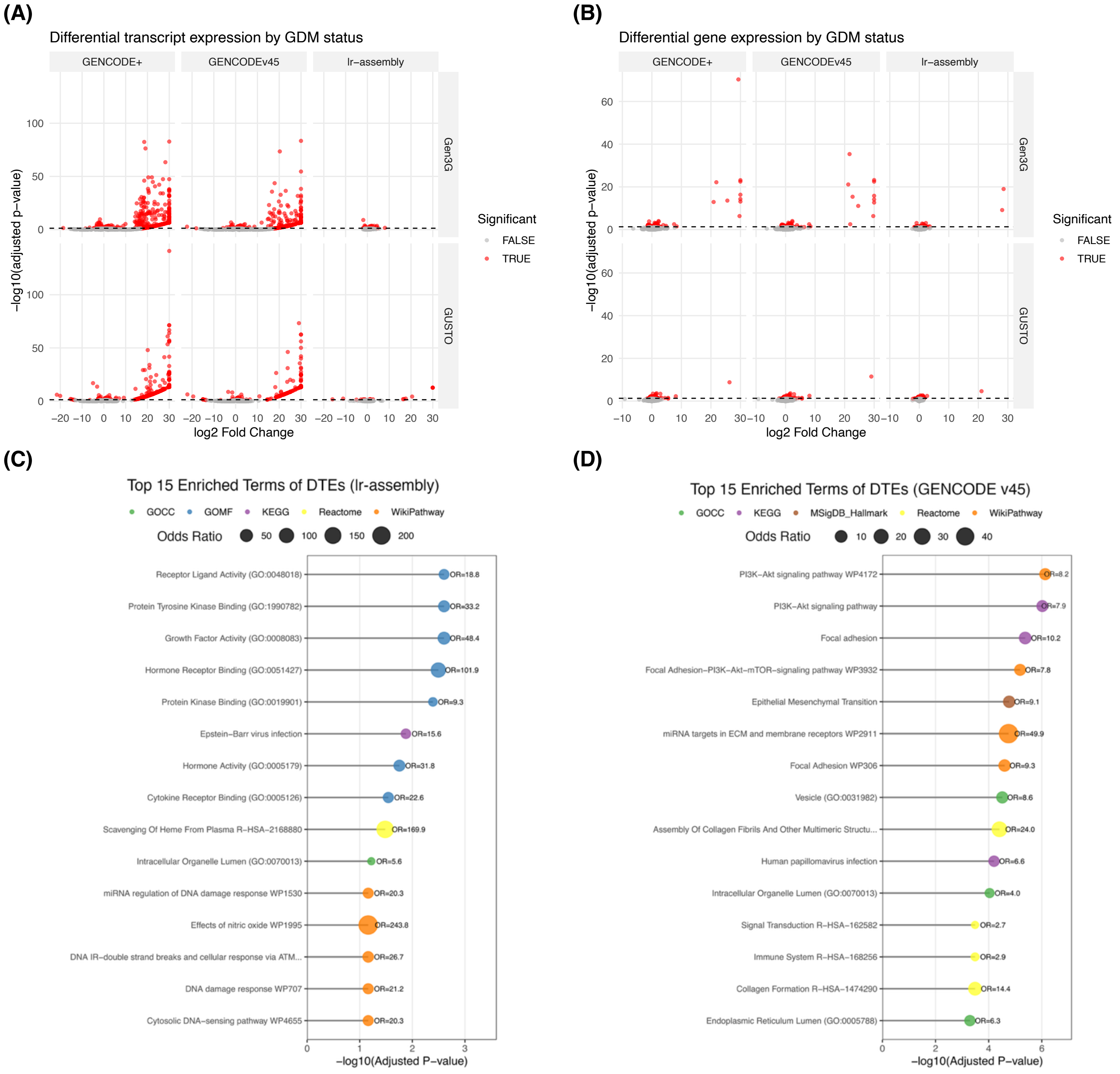


**Figure S7** – **Differential expression analyses** in placenta by GDM status at the **(A)** transcript and **(B)** gene levels, for both GUSTO and Gen3G given quantification against either the placenta transcriptome reference, GENCODEv45 or their union (GENCODE+). **(C-D)** Gene Ontology enrichment of GDM-associated transcripts in GUSTO given quantification against **(C)** the placenta transcript annotations and **(D)** GENCODEv45.

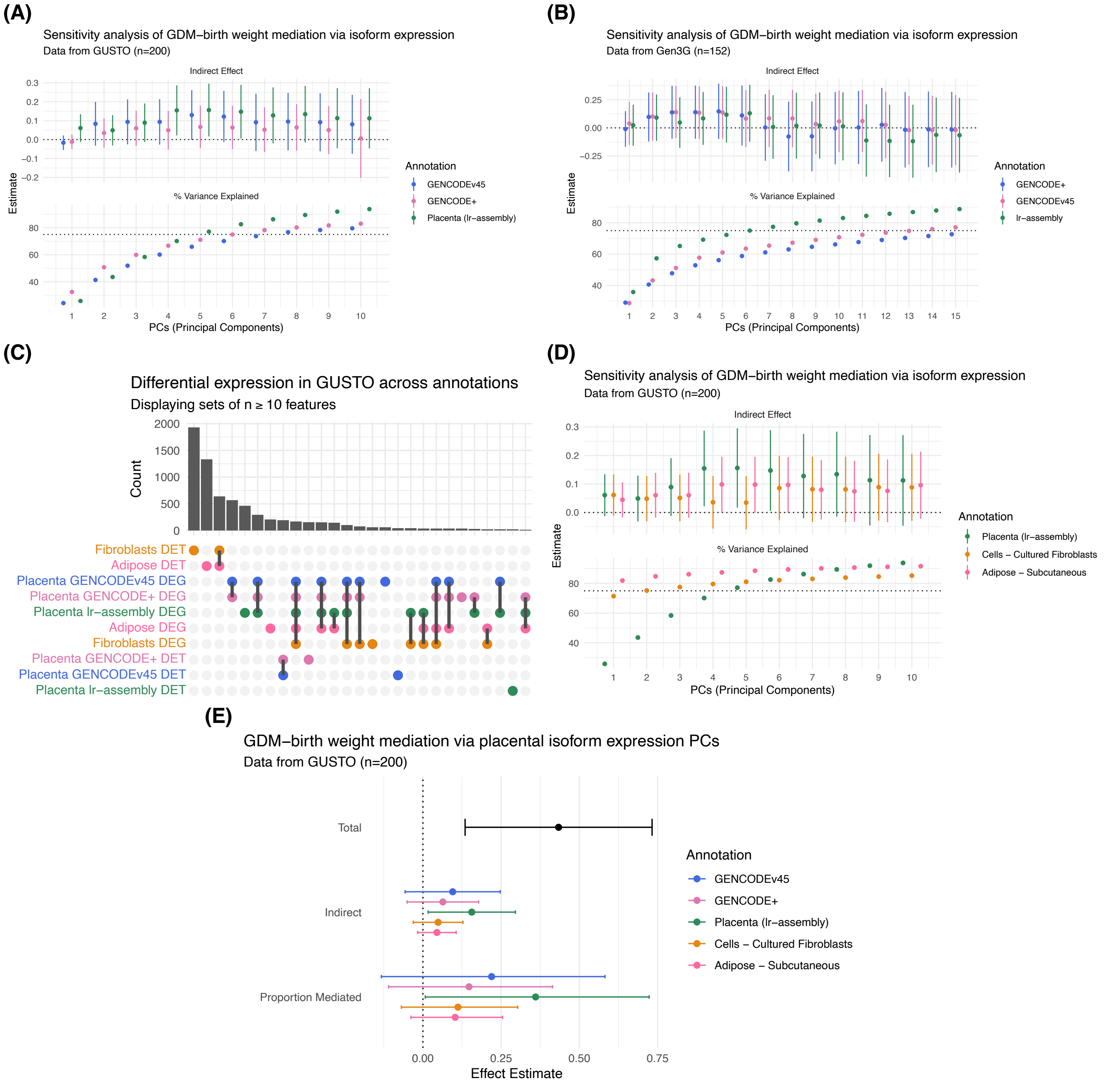


**Figure S8** – **Isoform expression causal mediation analyses. (A-B)** Sensitivity analysis of the indirect effect of placental transcript expression on GDM-birth weight associations in **(A)** GUSTO and **(B)** Gen3G when short-read samples were quantified against the placenta transcriptome reference, GENCODEv45 or their union (GENCODE+). Dashed lines at 0 for indirect effects and 75% for percent variance in isoform expression. **(C)** The count of differentially expressed genes (DEGs) and transcripts (DETs) in each cohort given short-read quantification against each annotation. **(D)** Sensitivity analysis of the indirect effect of placental transcript expression on GDM-birth weight associations in GUSTO when short-read samples were quantified against the placenta-, subcutaneous adipose-, or fibroblast-specific transcripts. Dashed lines at 0 for indirect effects and 75% for percent variance in isoform expression. **(E)** Mediation analysis effect estimates showing the relationship between gestational diabetes mellitus (GDM) and birth weight through placental isoform expression in GUSTO. Principal components were derived from differentially expressed transcripts for each annotation. Mediators included the first *PC* principal components that cumulatively explain at least 75% of expression variance.
